# Lung cancer-intrinsic SOX2 expression mediates resistance to checkpoint blockade therapy by inducing Treg-dependent CD8^+^ T cell exclusion

**DOI:** 10.1101/2023.09.06.556520

**Authors:** Elen Torres-Mejia, Sally Weng, Kim Nguyen, Ellen Duong, Leon Yim, Stefani Spranger

**Author notes:** Department of Molecular Biology, University of Texas Southwestern Medical Center, Dallas, TX, USA. Genentech, South San Francisco, CA 94080, USA. Corresponding author: Stefani Spranger, Koch Institute for Integrative Cancer Research at MIT, 77 Massachusetts Avenue, 76-453. Cambridge, MA 02139, USA.

## Abstract

Tumor-intrinsic signaling pathways can drastically affect the tumor immune microenvironment (TME), promoting tumor progression and resistance to immunotherapy by excluding immune cell populations from the tumor. Several tumor-cell intrinsic pathways have been reported to affect myeloid cell infiltration and downstream T cell infiltration. Clinical evidence suggests that the exclusion of cytotoxic T cells from the tumor core likewise mediates resistance. Here, we find that tumor cell-intrinsic SOX2 expression induces the exclusion of cytotoxic T cells from the tumor core and promotes resistance to checkpoint blockade therapy. CD8^+^ T cell exclusion was dependent on regulatory T cell-mediated suppression of tumor vasculature. Depleting tumor-infiltrating regulatory T cells via Glucocorticoid-Induced TNFR-Related (GITR) restored CD8^+^ T cell infiltration and reduced tumor growth in combination with checkpoint blockade therapy.

**Significance:** We identified tumor cell-intrinsic SOX2 expression in lung cancer as a mechanism of resistance to immunotherapy. SOX2 expression increases regulatory T cell populations in the TME, negatively affecting the tumor vasculature and blunting CD8^+^ T cell infiltration into the tumor core. This effect could be reverted by targeting regulatory T cells with anti-GITR therapy.

## Introduction

Immunotherapies, such as checkpoint blockade therapy (CBT), have significantly improved the survival of cancer patients^1^. Clinical studies in non-small cell lung cancer (NSCLC) have shown that combination of anti-PD-1 with anti-CTLA-4 significantly improves patients’ overall survival compared to patients treated only with chemotherapy^2^. However, only 30% of patients respond to the treatment, leaving 70% with no or only a modest overall survival extension. Several studies have shown that a pre-existing cytotoxic CD8^+^ T cell infiltrate in the tumor microenvironment (TME) is highly predictive of response to CBT^3,4^. Specifically in NSCLC, clinical data suggest the presence of four distinct tumor immune microenvironments: (a) T cell-infiltrated tumor with PD-L1 expression, (b) T cell-infiltrated tumor lacking PD-L1 expression; clinically referred to as non-functional T cell response (c), tumor lacking immune cell infiltration including CD8^+^ T cells and (d) T cell-excluded tumors, characterized by the presence of CD8^+^ T cells at the invasive margin of the tumor^5,6^. Data from clinical trials suggest that the T cell-infiltrated and PD-L1 positive tumors are most likely to respond to CBT treatment^6^. Understanding the mechanisms that lead to these different tumor immune microenvironments will facilitate the development of rational combination treatment strategies for NSCLC patients. It is conceivable that in particular T cell-excluded tumors, might be highly receptive to combination therapy, as a tumor-reactive T cell response is present yet fails to infiltrate the tumor core. However, to date the mechanisms mediating immune T cell exclusion in lung cancer are poorly understood.

Tumor cell-intrinsic signaling pathways can modulate the immune response during tumor progression and thus impact the tumor immune microenvironment. For example, *KRAS* and *LKB1/STK11* mutations in lung cancer have been associated with immune deserts, suggesting that tumor cell-intrinsic signaling pathways can directly impact T cell infiltration^7,8^. In NSCLC, increased expression of SOX2 is detected in approximately 20-65% of lung squamous cell carcinomas (LUSC) and 6-20% of lung adenocarcinomas (LUAC), the two major subtypes of NSCLC^9,10^. Although only a small fraction of LUAC patients exhibit high SOX2 expression, detection of SOX2 at the protein level indicates a poor prognosis on stage I of lung adenocarcinoma^11^. Furthermore, previous work demonstrated that SOX2 in LUSC cancer cells regulates the expression of the chemokine CXCL5 which promotes the recruitment of immune inhibitory neutrophils, favoring squamous tumorogenesis^12^. While the role of SOX2 in tumor progression has been extensively studied^13^, the mechanisms by which SOX2 affects T cell infiltration require more investigation.

In this study, we analyzed data from The Cancer Genome Atlas (TCGA) and biopsies from NSCLC patients and found that SOX2 expression correlates with CD8^+^ T cell exclusion from the tumor core. To determine how tumor-cell intrinsic activation of SOX2 changes the TME and evade immune surveillance, we established a pre-clinical model of SOX2 overexpression in lung cancer. We found that SOX2 induces CD8^+^ T cell exclusion from the tumor core and promotes resistance to CBT. CD8^+^ T cell exclusion was associated with increased density of regulatory T (Treg) cells in the peritumoral region of SOX2^+^ tumors. Mechanistically, Treg cells blunted activation of the tumor vasculature, preventing CD8^+^ T cells from entering the tumor core. Depletion of tumor Treg cells by anti-GITR^+^ treatment significantly improved CD8^+^ T cell infiltration in the tumor core and restored the responsiveness to CBT. In sum, we demonstrated that tumor cell-intrinsic expression of SOX2 mediates immune evasion by excluding CD8^+^ T cells from the tumor core via recruitment of Treg cells to the peritumoral regions in NSCLC.

## Results

### Tumor cell-intrinsic expression of SOX2 mediates T cell exclusion in NSCLC

To unbiasedly identify tumor-intrinsic signaling pathways that correlate with a non-T cell infiltrated phenotype in NSCLC, we segregated data from 999 NSCLC patients (LUSC and LUAC) from TCGA according to a previously published T cell-inflamed signature^14^. The NSCLC data set stratified in 636 (63.7%) patients predicted to have a high degree of T cell infiltration and 136 (13.6%) patients predicted to have a low degree of T cell infiltration. 227 (22.7%) could not be confidentially assigned and were categorized as a medium cohort (Fig. 1A). We then performed a pathway analysis using the differentially expressed genes between the predicted T cell high and T cell low patient cohorts. This analysis suggested a negative correlation between the SOX2 signaling pathway and T cell infiltration. Specifically assessing for *SOX2* expression levels in all T cell high, medium, and low cohorts showed that *SOX2* mRNA was indeed highest expressed in samples with low T cell infiltration (Fig. 1B). To validate whether tumor cell-intrinsic expression of SOX2 on the protein level is indeed associated with low T cell infiltration, we analyzed biopsies from LUSC and LUAC patients (Table S1) by multiplex immunofluorescence imaging (mIF) (Fig. 1C, 1D, and Fig. S1A). We determined SOX2^+^ tumor cell density, SOX2 mean fluorescence intensity (MFI), and cytotoxic CD8^+^ T cell density for each region independently. We observed that SOX2 protein levels were significantly higher in LUSC compared to LUAC (LUSC MFI = 24.28 vs. LUAC MFI = 4.002; Fig. S1B). Similarly, we found a higher frequency of SOX2-expressing cells in LUSC compared to LUAC tumor regions (91.6% in LUSC vs. 33.3% of LUAC, Table S1), as previously reported^10,13,15^. Interestingly, CD8^+^ T cell density was lower in LUSC compared to LUAD (142.3 cells/mm^2^ ± 20.19 SEM in LUSC vs. 196.5 cells/mm^2^ ± 16.42 SEM in LUAC; Fig. S1C). To determine whether a lack of T cell infiltration correlates with increased SOX2 expression in tumor cells, we classified each tumor region into low-T cell infiltrated (0-100 T cells/mm^2^) or high-T cell-infiltrated (100-400 T cells/mm^2^). Strikingly, we identified that tumor regions with low T cell infiltration had a higher density of SOX2^+^ tumor cells (Odds ratio=3.381; p<0.0001). In contrast, tumor regions with increased T cell infiltration were predominantly SOX2-negative or had low SOX2 expression (Fig. 1C and D). Detailed analysis of the tumor margin revealed that SOX2^+^ tumors were not entirely deprived of CD8^+^ T cell infiltration but that CD8^+^ T cells accumulated in the peritumoral region (Fig. 1D). These results suggested that in NSCLC, tumor cell-intrinsic overexpression of SOX2 acts as an immune evasion mechanism by inducing exclusion of T cells from the TME.

**Figure 1:**
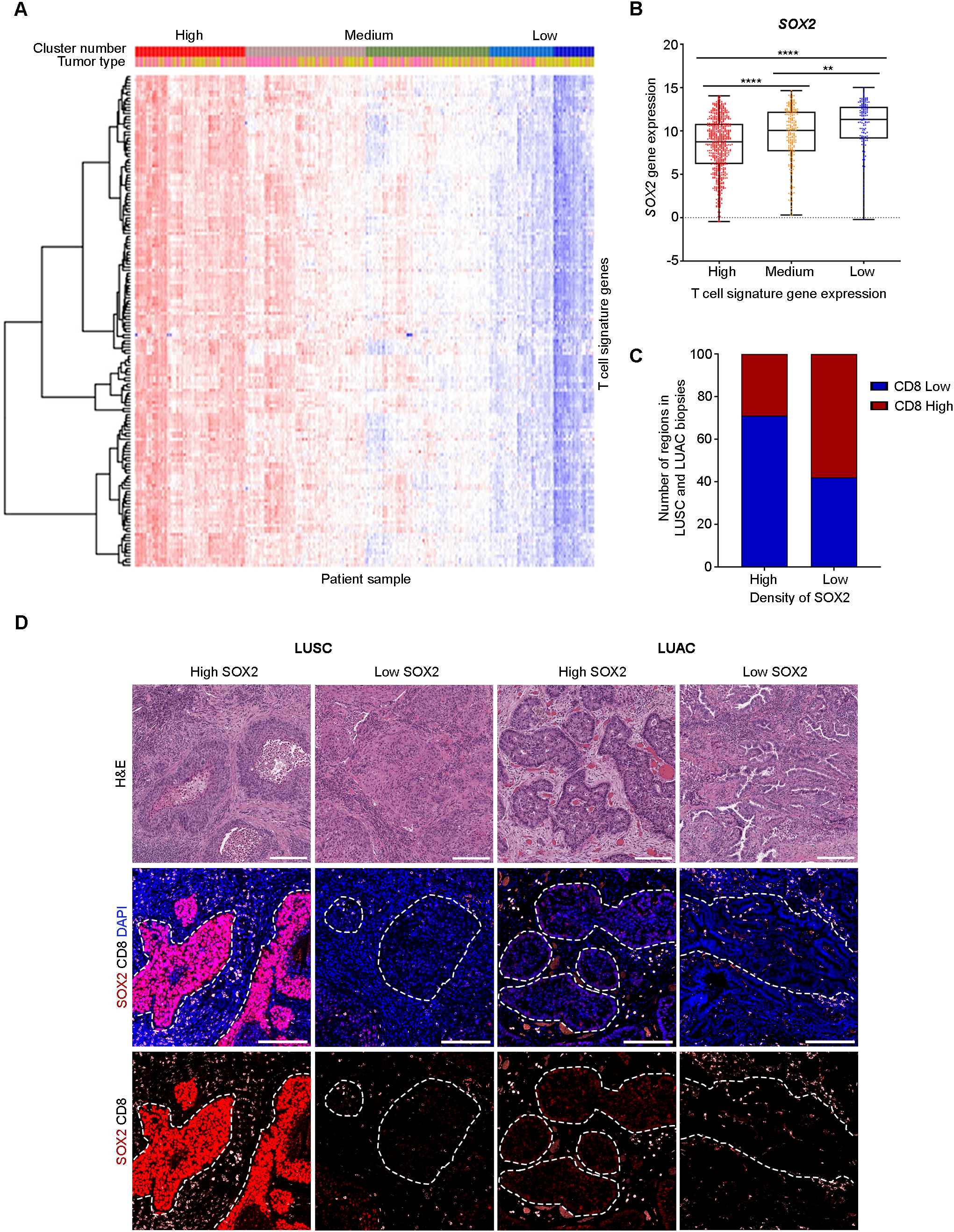
SOX2 expression in NSCLC correlates with low T cell infiltration. **A,** Heatmap shows segregation of NSCLC patient data from TCGA according to the expression of 160 T cell signature genes. **B,** Expression of *SOX2* in NSCLC patients with low, medium, and high expression levels of the T cell signature genes. Boxplot shows the median with minimum and maximum values. p-value was determined by one-way ANOVA. **C,** Quantification of CD8^+^ TILs in SOX2^+^ and SOX2^-^ regions in biopsies from LUSC and LUAD patients (257 tumor regions). Graph shows the number of defined regions with high (red) and low CD8^+^ TILs (blue) in low and high SOX2 density tumor regions. p-value was determined by Fisher’s exact test; p<0.0001; odd ratio=3.381. **D,** Representative H&E and fluorescence images showing immunostaining of CD8 (White) and SOX2 (Red) in biopsies from LUSC and LUAC patients. Nuclei were counterstained with DAPI (Blue). Dashed line indicates tumor regions; scale bar = 150 μm. When p-value is significant, results are flagged with stars, ****p<0.0001; ***p<0.0005; **p<0.005.

To understand the immune-modulatory role of SOX2 independently of its known function in LUSC differentiation^16^, we established a syngeneic mouse model of lung cancer using the mouse lung adenocarcinoma cell line KP driven by *Kras^G12D^* and *Trp53^-/-^* deletion^17^ (Fig. S2A). We evaluated SOX2 protein levels in the KP cell line by flow cytometry and found highly heterogeneous expression levels of SOX2 (Fig. S2B). To circumvent this heterogeneity, we established clonal cell lines with low levels of SOX2 expression. We used a clonally derived cell line (KP4C) to generate SOX2 overexpressing (KPS2) or control (KPCt) cells by lentiviral transduction. The generated cell lines were subsequently selected for high levels of SOX2 and low expression of SOX2, KPS2^High,^ and KPCt^Low^, respectively (Fig. S2A, S2B).

To obtain initial insights into the tumor immune microenvironment, we inoculated KPS2 and KPCt tumor cells subcutaneously in the flank of C57BL/6 mice. KPS2 tumor growth was significantly faster compared to KPCt tumors (Fig. 2A) in immunocompetent wild-type (WT) mice. In contrast, KPS2 and KPCt tumors grow similarly in immunocompromised Rag2^-/-^ mice, suggesting that the adaptive immune response mediates some level of control against KPCt but not KPS2 tumors (Fig. 2B). This phenotype was amplified when we used KPS2^High^ and KPCt^Low^ tumor cell lines, as KPCt^Low^ tumors were spontaneously cleared in WT mice but not in Rag2^-/-^ mice (Fig. S2C), while KPS2^High^ tumors grew progressively in both mouse models (Fig. S2D). To evaluate whether similar kinetics can also be observed for tumors forming in the lung, we engrafted KPS2 and KPCt orthotopically (Fig. 2C). We did not observe significant differences in lung tumor burden between these two lines in WT mice at day 21 post-tumor induction (Fig. 2C), which is consistent with our previous observation that T cell responses towards orthotopic KP tumors fail to control tumor growth^18^. Taken together, these results show that tumor cell-intrinsic upregulation of SOX2 mediates immune evasion from anti-tumor immunity.

**Figure 2:**
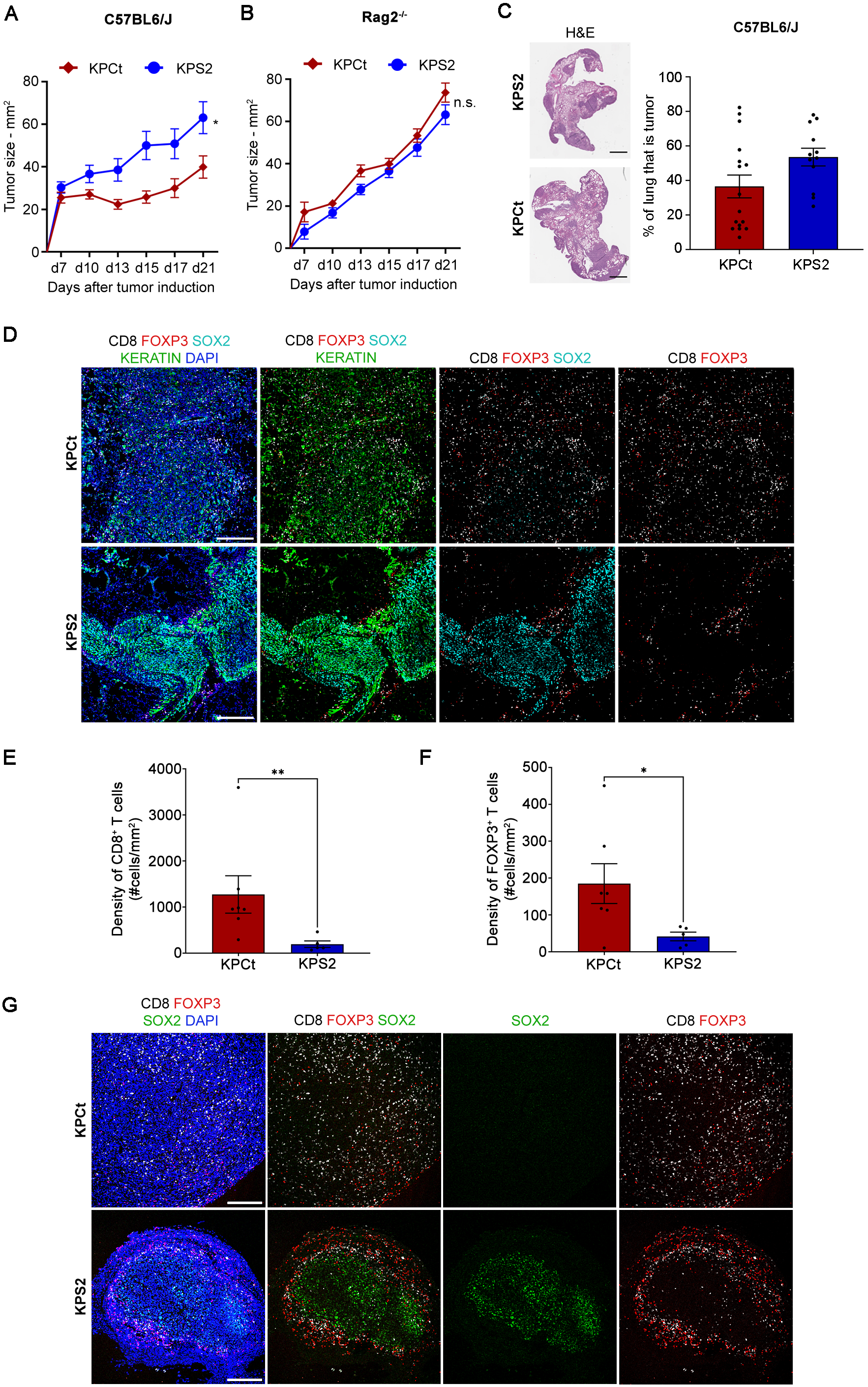
SOX2 overexpression in lung adenocarcinoma cells increases tumor growth and induces T cell exclusion from the tumor core. **A and B,** Outgrowth of KPCt and KPS2 flank tumors in **(A)** C56BL/6 and **(B)** Rag2^-/-^ mice. p-value at day 21 was determined by Mann-Whitney test (N=9). **C,** Representative H&E and lung tumor burden calculation at day 21 after tumor implantation with KPCt and KPS2 cell lines in C56BL/6 mice. Scale bar = 1000 μm. p-value at day 21 was determined by Mann-Whitney test (N=12-17). **D,** Representative fluorescence images and quantification of SOX2 (red), CD45 (White), and panKeratin (green) in KPCt and KPS2 lung tumors at day 21. Nuclei were counterstained with DAPI (Blue). Scale bar = 300 μm. **E and F,** Quantification of CD8^+^ and FOXP3^+^ T cell densities in KPCt and KPS2 flank tumors (N=5-7). **G,** Representative confocal images of SOX2 (green), CD8 (white), and FOXP3 (red) in KPCt and KPS2 flank tumors at day 21 post-tumor implantation. Nuclei were counterstained with DAPI (Blue). Scale bar = 250 μm. p-value was determined by Mann-Whitney test. Data are shown as means ± SEM. When p-value is significant, results are flagged with stars, **p<0.005; *p<0.05.

Given the observed control of flank KPCt tumors in WT but not Rag2^-/-^ mice, we next characterized the tumor-infiltrating T cell populations in KPS2 and KPCt tumors by flow cytometry. We detected no differences in the percentage of CD8^+^ T cells, CD4^+^ T cells, and FOXP3^+^ Treg cells between the tumor conditions (Fig. S2E, S2F). Similarly, we did not observe differences in the total number of T cells between KPCt and KPS2 flank tumors (Fig. S2G). Expression of the T cell activation marker Programmed Cell Death Protein 1 (PD-1) was also similar between KPS2 and KPCt T cell infiltrates, suggesting a similar degree of antigen encounter (Fig. S2H). We made similar observations when analyzing the immune infiltrate of orthotopic KPCt and KPS2 lung tumors (Fig. S3A, S3B). Since we observed a T cell exclusion phenotype in human NSCLC samples (Fig. 1C-D), we next analyzed the spatial location of cytotoxic T cells (CD8^+^) and Treg cells (FOXP3^+^) in the TME by mIF. Consistent with the clinical observation, we observed low infiltration of T cells into the tumor core of orthotopic KPS2 lung tumors (CD8^+^ T cells = 194 cells/mm^2^ ± 71.06 SEM; Treg cells = 41.56 cells/mm^2^ ± 11.74 SEM), while the peritumoral region showed a profound presence of T cells (Fig. 2D-F). In contrast, KPCt tumors showed brisk infiltration of T cells (CD8^+^ T cells = 1273 cells/mm^2^ ± 406.4 SEM; Treg cells = 185 cells/mm^2^ ± 54 SEM) within the tumor core (Fig. 2D-2F). This phenotype was maintained in flank tumors, where we observed an accumulation of CD8^+^ T cells and Treg cells in the peritumoral region and low infiltration of both T cell populations in the core of KPS2 flank tumors (Fig. 2G). In contrast, KPCt flank tumors had a high infiltration of T cells in the tumor core (Fig. 2G). These results confirm our previous observation in NSCLC biopsies where SOX2 was associated with the exclusion of CD8^+^ T cells from the tumor core (Fig. 1C and 1D).

To determine whether SOX2-mediated exclusion was specific for T cells, we assessed overall immune infiltration using CD45 staining. Interestingly, we observed that KPS2 tumors were less infiltrated with CD45^+^ leukocytes than KPCt (CD45^+^ cells in KPCt = 7798 cells/mm^2^ ± 655.4 SEM vs. CD45^+^ cells in KPS2 = 3864 cells/mm^2^ ± 541.7 SEM; Fig. S3C, S3D). Previous studies indicated that a low number of dendritic cells (DC) within the tumor mediated a lack of T cell infiltration^19^. Thus, we assessed the density of DC by mIF, using the marker CD11c. Strikingly, there were no significant differences in the density of DC in both lung tumor conditions (CD11c^+^ cells in KPCt = 1593 cells/mm^2^ ± 311.7 SEM vs. CD11c^+^ cells in KPS2 = 1351 cells/mm^2^ ± 265.6 SEM; Fig. S3E, S3F). These results suggest that tumor cell-intrinsic SOX2 expression modifies the TME, impairing the infiltration of T cells.

### SOX2 induces resistance to checkpoint blockade therapy

Pre-clinical and clinical data in melanoma and NSCLC have shown that the pre-existence of cytotoxic T cells in the tumor core but not the peritumoral site correlates with a positive response to CBT^4,5,19–21^. To evaluate whether T cell exclusion, mediated by tumor cell-intrinsic SOX2, impacts the response to CBT, we treated KPCt and KPS2 flank tumor-bearing mice with dual CBT, anti-PD-L1 plus anti-CTLA-4 (Fig. 3A). As we have previously shown that orthotopic KP lung tumors fail to respond to CBT, due to the acquisition of a dysfunctional and tolerogenic T cell phenotype^18^; we focused our analysis on flank tumors. CBT treatment induced tumor regression and stable disease in KPCt tumors (Fig. 3B). Strikingly however, CBT treatment did not yield any therapeutic benefit in KPS2 tumors resulting in progressive growth similar to untreated tumors (Fig. 3B). We next evaluated whether CBT treatment affected the infiltration of cytotoxic T cells and Treg cells by mIF and observed that the density of both T cell populations doubled after CBT treatment in KPCt flank tumors, consistent with previous reports^18^ (Fig. 3C, 3D). However, in KPS2, neither CD8^+^ T cells nor Treg cell density was increased (Fig. 3C, 3D). These results suggest that tumor cell-intrinsic SOX2 signaling mediates resistance to CBT by facilitating T cell exclusion.

**Figure 3:**
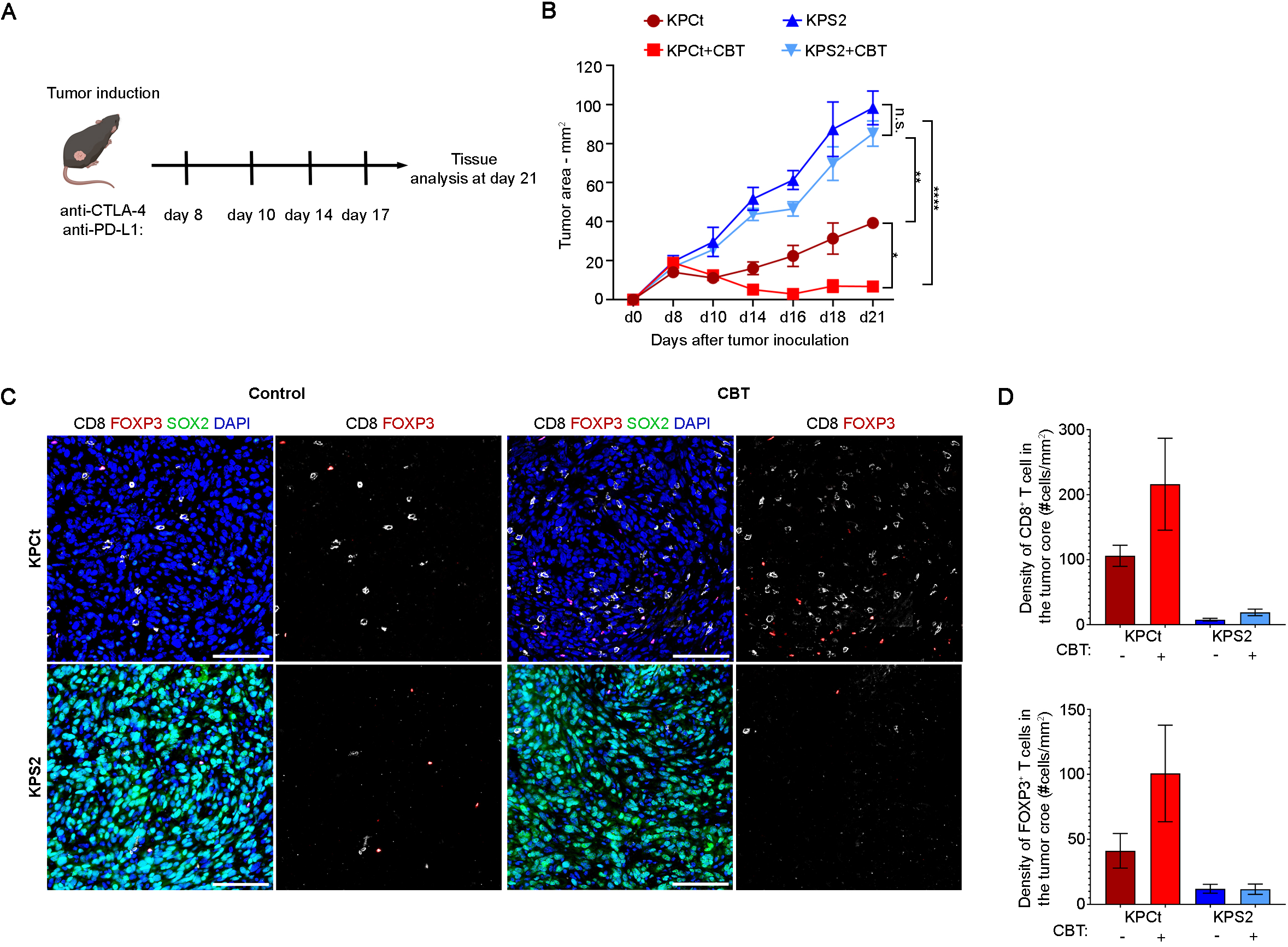
SOX2^+^ flank tumors are resistance to CBT. **A,** Experimental design for (B-D). **B,** Outgrowth of flank KPCt and KPS2 tumors treated with CBT or control. p-value at day 21 was determined by One-way ANOVA (N=5). **C)** Representative fluorescence images showing immunostaining of SOX2 (Green), CD8 (White), and FOXP3 (Red) in KPCt and KPS2 flank tumors at day 21 treated with CBT or control. Nuclei were counterstained with DAPI (Blue). Scale bar = 100 μm. **D)** Quantification of CD8^+^ and FOXP3^+^ T cell densities in KPCt and KPS2 flank tumors at day 21 treated with CBT or control. p-value was determined by One-way ANOVA. Data are shown as means ± SEM. When p-value is significant, results are flagged with stars, ****p<0.00005; **p<0.005; *p<0.05.

### Tumor cell-intrinsic SOX2 expression does not affect T cell priming and systemic immunity but blunts effector T cell infiltration into the tumor core

To interrogate the activation and trafficking of tumor-reactive CD8^+^ T cells, we engineered the KPS2 and KPCt cell lines to express the model antigen SIYRYYGL (SIY) fused to GFP. KPS2.SIY and KPCt.SIY cells were sorted for similar GFP expression levels (Fig. S4A). We first evaluated the effect of SIY expression on tumor outgrowth by inducing flank and orthotopic lung tumors with KPS2.SIY and KPCt.SIY cells. Strikingly, we observed that KPCt.SIY flank tumors started to regress in a T cell-dependent manner (Fig. 4A). Consistent with our previous observation that tumor cell-intrinsic SOX2 signaling mediates immune evasion, we observed that KPS2.SIY flank tumors grew similarly in WT and Rag2^-/-^ mice (Fig. 4B). We validated this observation using KPS2.SIY and KPCt.SIY orthotopic lung tumors and observed similar tumor burden between WT and Rag2^-/-^ mice for KPS2.SIY tumors while KPCt.SIY showed 69.2% of tumor control (Fig. 4C-4D). It is possible that tumor cell-intrinsic SOX2 overexpression confers resistance to CD8^+^ T cell-mediated cytotoxicity. To test this possibility, we performed an *in vitro* killing assay using *ex vivo* activated SIY-specific TCR-transgenic CD8^+^ T cells (2C T cells) co-cultured with tumor cells. Comparing targeted lysis of KPCt.SIY or KPS2.SIY over a range of effector-to-target ratios did not show any significant differences in cytotoxicity (Fig. 4E), suggesting that both cell lines are equally susceptible to CD8^+^ T cell-mediated lysis. To ensure that the addition of the model antigen SIY does not affect the functional state of tumor-specific T cells, we phenotypically analyzed SIY-reactive T cells in subcutaneous tumors at day 8 and lung tumors at day 21 using flow cytometry. There were no significant differences in PD-1 or Granzyme B (GzmB) expression by the endogenous SIY^+^ reactive T cells in the flank tumor (Fig. S4B-S4D). The only notable difference was an increase in the percentage of SIY^+^ reactive T cells in KPS2 tumors compared to KPCt flank tumors (Fig. S4B). A reduction in activation-induced cell death may explain this increase in tumor-reactive T cells in the TME of KPS2 tumor, a phenomenon often observed in T cell-infiltrated TMEs^22^. In the orthotopic lung tumors, consistent with the observations made using the KPCt and KPS2 parental cell lines (Fig. S3A, S3B), we found similar frequencies of SIY^+^ reactive T cells and PD-1 expression (Fig. S4E, S4F). Taken together, these results indicate that tumor cell-intrinsic SOX2 signaling mediates immune evasion even in the presence of a strong antigen.

**Figure 4:**
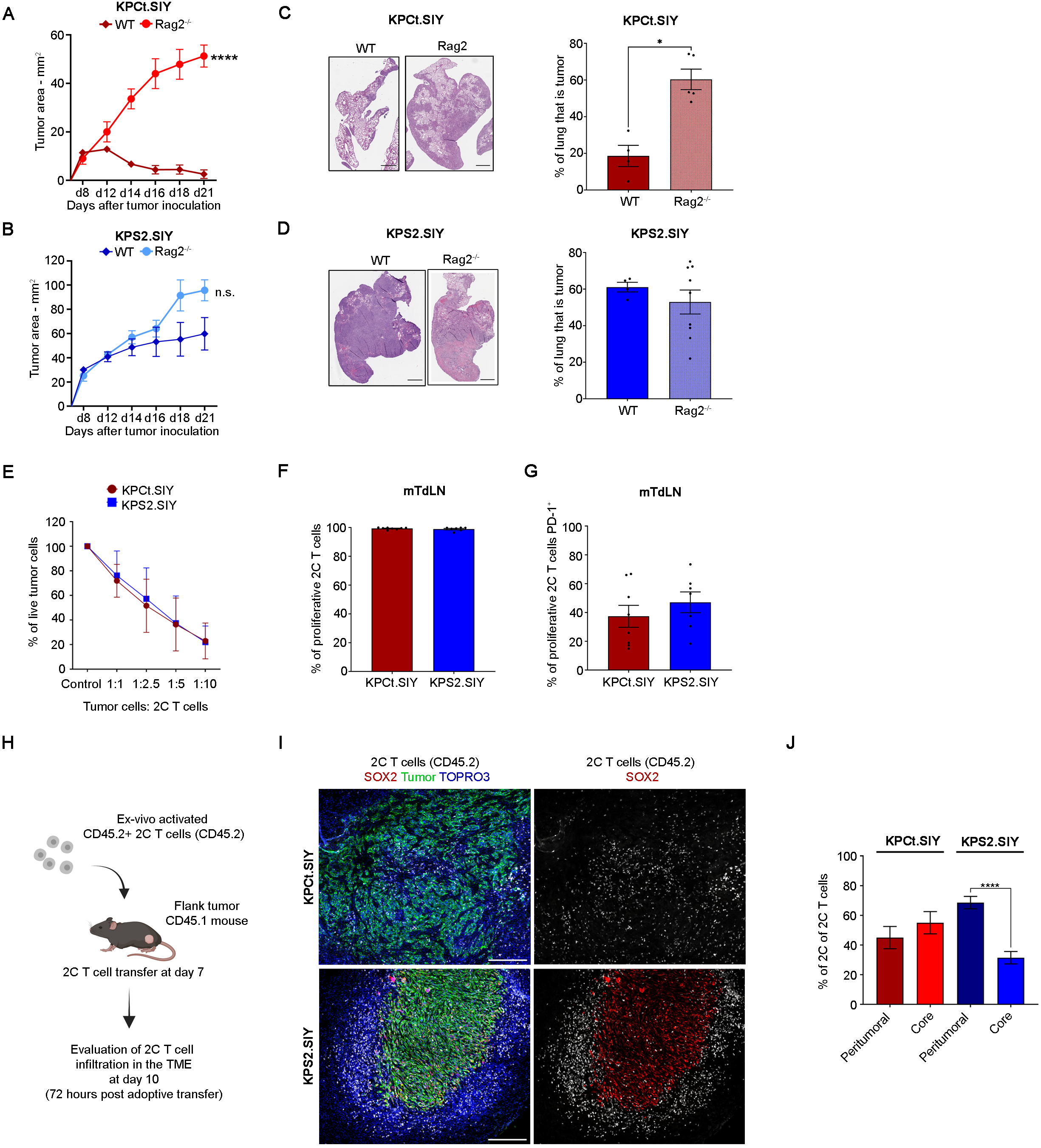
SIY-specific T cells are recruited to the tumor site but failed to infiltrate SOX2^+^ tumors. **A and B,** Outgrowth of KPCt.SIY (A) and KPS2.SIY (B) flank tumors in WT (C56BL/6) and Rag2^-/-^ mice. p-value at day 21 was determined by Mann-Whitney test (N=5-9). **C and D,** Representative H&E images and lung tumor burden quantification in WT (C56BL/6) and Rag2^-/-^ mice at day 21 post-tumor induction with KPCt.SIY (C) or KPS2.SIY (D) cell lines. p-value was determined by Mann-Whitney test (N=4-9). **E,** Cell survival of KPCt.SIY and KPS2.SIY co-cultured with *ex-vivo* activated 2C T cells at different tumor to 2C T cell ratios. Cell survival was measured by the percentage of GFP positive cells at 17 hours post co-culture with 2C T cells. p-value was determined by two-ways ANOVA (N=3, experiment was repeated three times). **F and G,** Quantification of the percentage of (F) CFSE-labeled 2C T cells that proliferate and (G) PD-1^+^ 2C T cells in the mTdLN of KPCt.SIY and KPS2.SIY lung-tumor bearing mice at 72 hours after adoptive transfer, day 10 of tumor growth. p-value was determined by t-student (N=7-8). **H,** Experimental design for Fig. 4I and 4J. **I and J,** Representative fluorescence images and quantification of KPCt.SIY and KPS2.SIY flank tumors at 72 hours after adoptive transfer of ex-vivo activated 2C T cells (CD45.2) into WT (C56BL/6) mice (CD45.1), day 10 of tumor growth. (I) Images show GFP expression by tumor cells (Green), CD45.2 (White), and SOX2 (Red). Nuclei were counterstained with TO-PRO-3 (Blue). Scale bar = 250 μm. (J) Graph shows the percentage of 2C T cells in the tumor core and in the peritumoral region of flank tumors. p-value was determined by t-student (N=5). Data are shown as means ± SEM. When p-value is significant, results are flagged with stars, ****p<0.0001, *p<0.05.

Previous studies have shown that activation of tumor cell-intrinsic oncogenic pathways can impair T cell priming, leading to poor systemic immunity and as a consequence a low number of tumor-infiltrating CD8^+^ T cells ^19,23^. To determine whether the expression of SOX2 in tumor cells negatively affects CD8^+^ T cell priming, we transferred CFSE-labeled naïve 2C T cells into lung tumor-bearing mice at day seven post-tumor induction (Fig. S4G). Three days after adoptive T cell transfer, we collected the tumor-draining lymph node (mediastinal lymph node, mTdLN) to assess the proliferation (CFSE dye dilution) of the transferred 2C T cells as a readout for T cell activation (Fig. S4G). We observed a similar degree of 2C T cell proliferation between KPS2.SIY and KPCt.SIY in the mTdLN (Fig. 4F and Fig. S4H), suggesting that T cell priming is unaffected by tumor cell-intrinsic SOX2 overexpression. To affirm this notion, we assessed PD-1 expression levels on the transferred 2C T cells in the mTdLN. We observed similar percentages of proliferative 2C T cells PD-1^+^ in the mTdLN of KPS2.SIY orthotopic lung tumors (Fig. 4G and Fig. S4H). To evaluate the functional properties of the activated effector CD8^+^ T cells, we assessed the peripheral immunity towards the tumors using IFNγ-ELISpot. Consistent with our observation that T cell priming is similar between both tumor settings, we did not detect significant differences in the number of endogenous SIY-reactive IFNγ-producing T cells in the spleens of mice with flank or orthotopic KPS2 or KPCt tumors (Fig. S4I, S4J). We did observed that the systemic immunity induced in response to orthotopic lung cancer was diminished compared to flank tumors, which is consistent with our previous report^18^. These results suggest that T cell priming and systemic antitumor immunity are unaffected by tumor cell-intrinsic SOX2 overexpression.

Given that we observed similar systemic T cell priming and systemic immunity in mice bearing KPCt.SIY and KPS2.SIY tumors, we next investigated the recruitment of effector CD8^+^ T cells to SOX2^+^ tumors. To separate T cell activation from T cell homing and infiltration, we performed an adoptive transfer using *ex vivo* activated effector 2C T cells (Fig. 4H). We analyzed the spatial localization of transferred 2C T cells using the congenic marker CD45.2. Strikingly, we observed that SOX2^+^ KPS2.SIY flank tumors excluded tumor-reactive effector 2C T cells from the tumor core (Fig. 4I). In contrast, 2C T cell homing to the tumor margin was unaffected (Fig. 4I). We quantified the percent of 2C T cells in the tumor core versus the peritumoral site and observed that 69% of T cells were located in the peritumoral area of KPS2.SIY flank tumors (Fig. 4J). In contrast, KPCt control tumors showed brisk infiltration with 2C T cells, with a slight increase of infiltration in the tumor core (55% tumor core; Fig. 4I-J). These findings demonstrate that tumor cell-intrinsic SOX2 signaling mediates T cell exclusion from the tumor core, while T cell priming and homing to the inflammatory site are unaffected.

### Regulatory T cells in the TME dampen CD8^+^ T cell infiltration

Our results show that tumor cell-intrinsic expression of SOX2 induces the exclusion of effector CD8^+^ T cells from the tumor core (Fig. 4I, 4J). We have previously observed that Treg cells accumulate in the peritumoral region of KPS2 tumors (Fig. 2G). Thus, we sought to further investigate the Treg cell distribution in the TME of flank tumors generated with the KPCt.SIY and KPS2.SIY cell lines. Strikingly, when we looked at day 10 post-tumor cell injection, we observed no differences in Treg density in the core (Fig. S5A, S5B). However, we found a significant accumulation of Treg cells in the peritumoral region of KPS2.SIY flank tumors compared to KPCt.SIY tumors (615.9 cells/mm^2^ ± 92.36 SEM in KPS2.SIY vs. 305.4 cells/mm^2^ ± 45.05 SEM in KPCt.SIY; Fig. S5A, S5C). Consistent with our findings for adoptively transferred effector 2C T cells (Fig. 4I, 4J), the infiltration of endogenous CD8^+^ T cells was significantly reduced in the core of KPS2.SIY tumors (1596 cells/mm^2^ ± 316.4 SEM in KPCt.SIY vs. 490.0 cells/mm^2^ ± 89.38 SEM in KPS2.SIY; Fig. S5A, S5D). Meanwhile, the peritumoral region showed a similar density of CD8^+^ T cells in KPCt.SIY and KPS2.SIY flank tumors (Fig. S5A, S5E). As Treg cells are immunosuppressive cells that can modulate the TME to dampen the antitumor immune response^24^, we hypothesized that their increased numbers in the periphery of KPS2 tumors could be responsible for the SOX2-mediated CD8^+^ T cell exclusion from the tumor core.

To assess whether Treg cells are required for the exclusion of CD8^+^ T cells from the tumor core, we induced KPS2.SIY and KPCt.SIY flank tumors in *FoxP3^DTR^* mice. Following tumor engraftment, Treg cells were depleted systemically by diphtheria toxin (DT) administration in KPS2.SIY tumor-bearing mice (Fig. 5A). We first assessed tumor growth in KPCt.SIY, KPS2.SIY, and KPS2.SIY+DT and found that KPS2.SIY flank tumors treated with DT were significantly smaller compared to KPS2.SIY untreated tumors. Specifically, KPS2.SIY+DT tumors showed a 54% reduction in tumor size, growing similarly to untreated KPCt.SIY flank tumors (Fig. S6A). We further confirmed that there was a 99% depletion efficiency of FOXP3^+^CD4^+^ Treg cells in the spleen after DT administration (Fig. S6B). We then used mIF to evaluate Treg cell depletion and CD8^+^ T cell infiltration in the TME. Following DT treatments, Treg cell density was significantly reduced in the TME of KPS2.SIY+DT flank tumors compared to untreated KPS2.SIY tumor-bearing mice, and this reduction was observed in the tumor core as well as in the peritumoral region (Fig. 5B-5D, Fig. S6C). Strikingly, we observed a significant increase in CD8^+^ T cell infiltration into the tumor core of KPS2.SIY DT-treated mice (417 cells/mm^2^ ± 76 SEM in KPS2.SIY vs. 1400 cells/mm^2^ ± 203.7 SEM in KPS2 DT-treated), reaching densities as high as the those observed in the core of KPCt.SIY flank tumors (1043 cells/mm^2^ ± 175.8 SEM in KPCt.SIY; Fig. 5E, and Fig. S6C). Furthermore, the peritumoral region of KPS2.SIY DT-treated tumors showed a significant increase in CD8^+^ T cell density compared to untreated and control tumors (Fig. 5F). We also evaluated the density of SOX2^+^ tumor cells and observed no changes after DT treatment, confirming that DT treatment did not affect SOX2 expression by the tumor cells *in vivo* (Fig. S6D). These results demonstrate that Treg cells in the KPS2 TME are sufficient to blunt the infiltration of effector CD8^+^ T cells into the tumor core.

**Figure 5:**
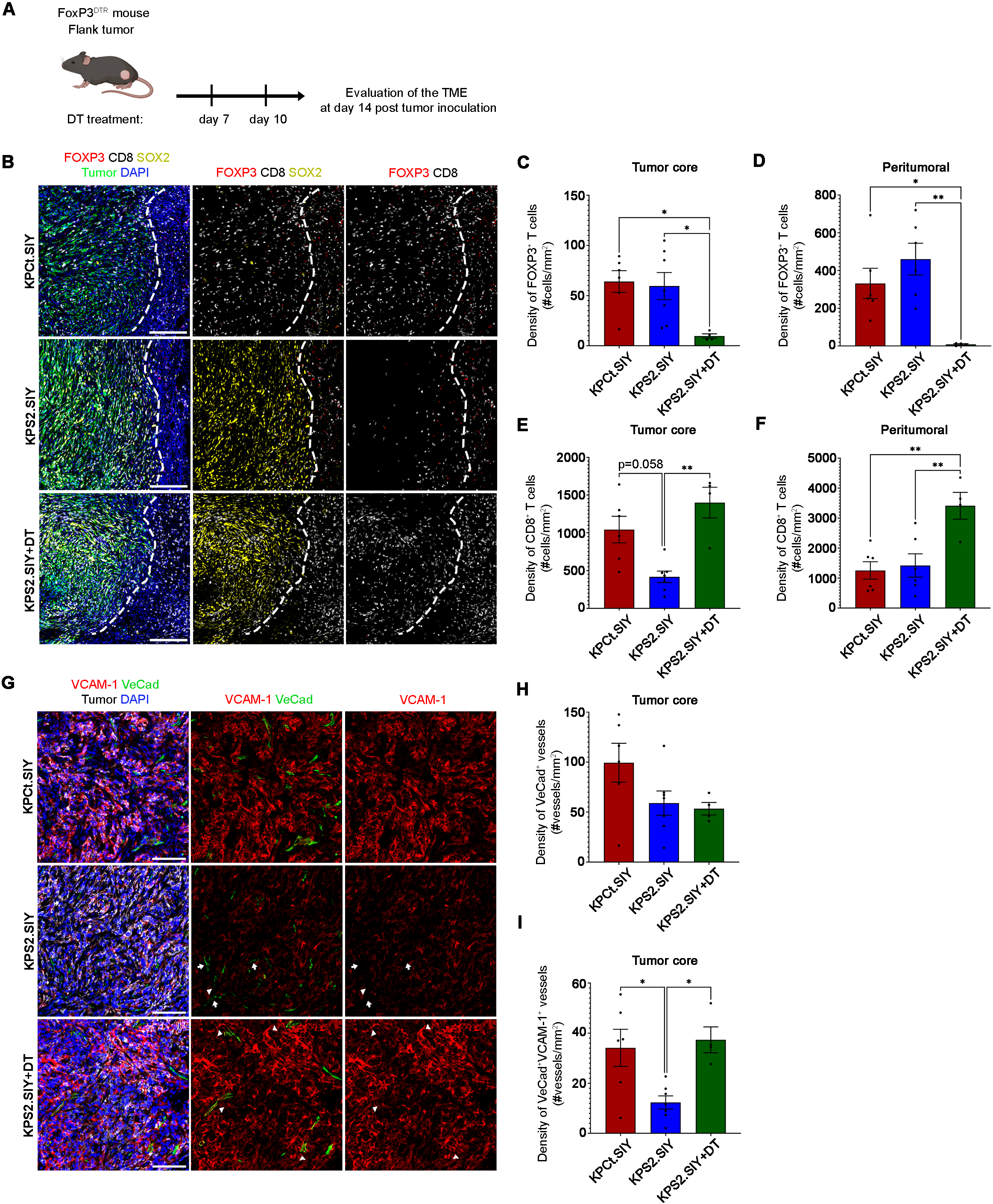
Depletion of FOXP3^+^ T cells in KPS2.SIY tumors improves CD8^+^ T cell infiltration and increases the expression of VCAM-1 in endothelial cells. **A,** Experimental design for (B-I). **B,** Representative fluorescence images of SOX2 (yellow), CD8 (white), FOXP3 (red), and tumor cells (green) in KPCt.SIY, KPS2.SIY, and KPS2.SIY+DT flank tumors at day 14 post-tumor induction. Nuclei were counterstained with DAPI (Blue). Scale bar = 100 μm. **C to F,** Quantification of FOXP3^+^ and CD8^+^ T cell densities in the core and peritumoral region of KPCt.SIY, KPS2.SIY, and KPS2.SIY+DT flank tumors. p-value was determined by one-way ANOVA (N=4-7). **G,** Representative fluorescence images of VeCAD (green), VCAM-1 (red), and tumor cells (white) in KPCt.SIY, KPS2.SIY and KPS2.SIY+DT flank tumors at day 14 post-tumor induction. Nuclei were counterstained with DAPI (Blue). Scale bar = 100 μm. **H and I,** Quantification of VeCAD^+^ vessel density (H) and VCAM-1^+^VeCAD^+^ vessel density (I) in the core of KPCt.SIY, KPS2.SIY, and KPS2.SIY+DT flank tumors. p-value was determined by one-way ANOVA (N=4-7). Data are shown as means ± SEM. When p-value is significant, results are flagged with stars, **p<0.01; *p<0.05.

While it has been demonstrated that Treg cells can directly affect the function and proliferation of CD8^+^ T cells^24^, thus far, less is known about their effect on CD8^+^ T cell migration. For effector CD8^+^ T cells to home into inflamed tissues, vascular endothelial cells need to acquire an activated state^25^. Activation of endothelial cells results in the upregulation of adhesion molecules that facilitate the rolling, adhesion, and extravasation of CD8^+^ T cells to the inflamed tissue^25^. Treg cells have been shown to promote tumor angiogenesis by expressing angiogenic factors, such as VEGF-A^26,27^. Tumor-associated vasculature is often unable to support T cell adhesion and extravasation^28^, and aberrant vasculature has been linked with poor T cell infiltration in humans and pre-clinical models^29,30^. Considering that the presence of peritumoral Treg cells in the TME of KPS2 tumors correlated with T cell exclusion (Fig. S5A-E), we hypothesized that Treg cells could impair the activation of endothelial cells, thus reducing T cell extravasation. To test this hypothesis, we analyzed the expression of the vascular cell adhesion molecule 1 (VCAM-1), a critical adhesion molecule necessary for CD8^+^ T cell extravasation^25^, in the TME following Treg cell depletion. We observed VCAM-1 expression in KPS2.SIY and KPCt.SIY tumor cells as well as in stroma cells (Fig. 5G). Analysis of KPS2.SIY and KPCt.SIY cells cultured *in vitro* confirmed the expression of VCAM-1 in both cell lines; however, KPS2.SIY cells showed a reduced expression level (Fig. S6E). We identified the tumor vasculature using the endothelial cell marker VE-cadherin (vascular endothelial cadherin, VeCad). We analyzed the vasculature in the core of KPS2.SIY flank tumors and determined the density of VCAM-1^+^ VeCad^+^ vessels in control and DT-treated mice (Fig. 5G and Fig. S6F). While there were no significant differences in vessel density between tumor conditions (Fig. 5G, 5H), we detected lower expression of VCAM-1 on vessels located in the core of KPS2.SIY flank tumors (Fig. 5G, 5I). Strikingly, however, after Treg cell depletion the number of VCAM-1^+^VeCad^+^ vessels in the tumor core significantly increased (12.34 vessels/mm^2^ ± 2.631 SEM in KPS2.SIY vs. 37.33 vessels/mm^2^ ± 5.159 SEM in KPS2.SIY DT-treated mice) to values similar to the KPCt.SIY tumors (34.14 vessels/mm^2^ ± 7.394 SEM; Fig. 5I). In sum, these data suggest that retention of Treg cells in the peritumoral region of SOX2^+^ tumors affects endothelial cell activation by preventing activation-associated upregulation of VCAM-1, thus blunting cytotoxic CD8^+^ T cell infiltration into the tumor core.

### Anti-GITR treatment reduces FOXP3^+^ Treg cells in the TME, and restores CD8^+^ T cell infiltration

Since systemic depletion of all Treg cells restores CD8^+^ T cell infiltration into the tumor core, we next aimed to identify a translationally relevant approach to specifically reduce Treg cell numbers in the TME. Depletion of Treg cells in the TME has been accomplished by others using the anti-GITR antibody, clone DTA-1. Glucocorticoid induced TNF receptor (GITR) is a member of the TNFR superfamily constitutively expressed at high levels in Treg cells^31,32^. Previous studies have shown that anti-GITR treatment in different cancer models significantly depletes Treg cells in the TME while increasing survival of CD8^+^ T cells^33–35^. In our case, GITR would be an ideal candidate for the targeted depletion of effector Treg cells in the TME. To determine whether targeting Treg cells in the TME with anti-GITR treatment improves CD8+ T cell infiltration, we treated flank tumor-bearing mice with the mouse anti-GITR antibody^36^ (Fig. 6A). We assessed changes in Treg cell and CD8^+^ T cell densities in the tumor core and peritumoral regions after anti-GITR treatment. We observed that anti-GITR treatment significantly reduced Treg cells in the core and in the peritumoral region of KPS2.SIY flank tumors (Fig. 6B-6D). Consistent with our previous results (Fig. 5), the reduction of Treg cells resulted in improved CD8^+^ T cell infiltration following anti-GITR treatment (1307 cells/mm^2^ ± 250.6 SEM in treated vs. 546.5 cells/mm^2^ ± 78.17 SEM untreated; Fig. 6E).

**Figure 6:**
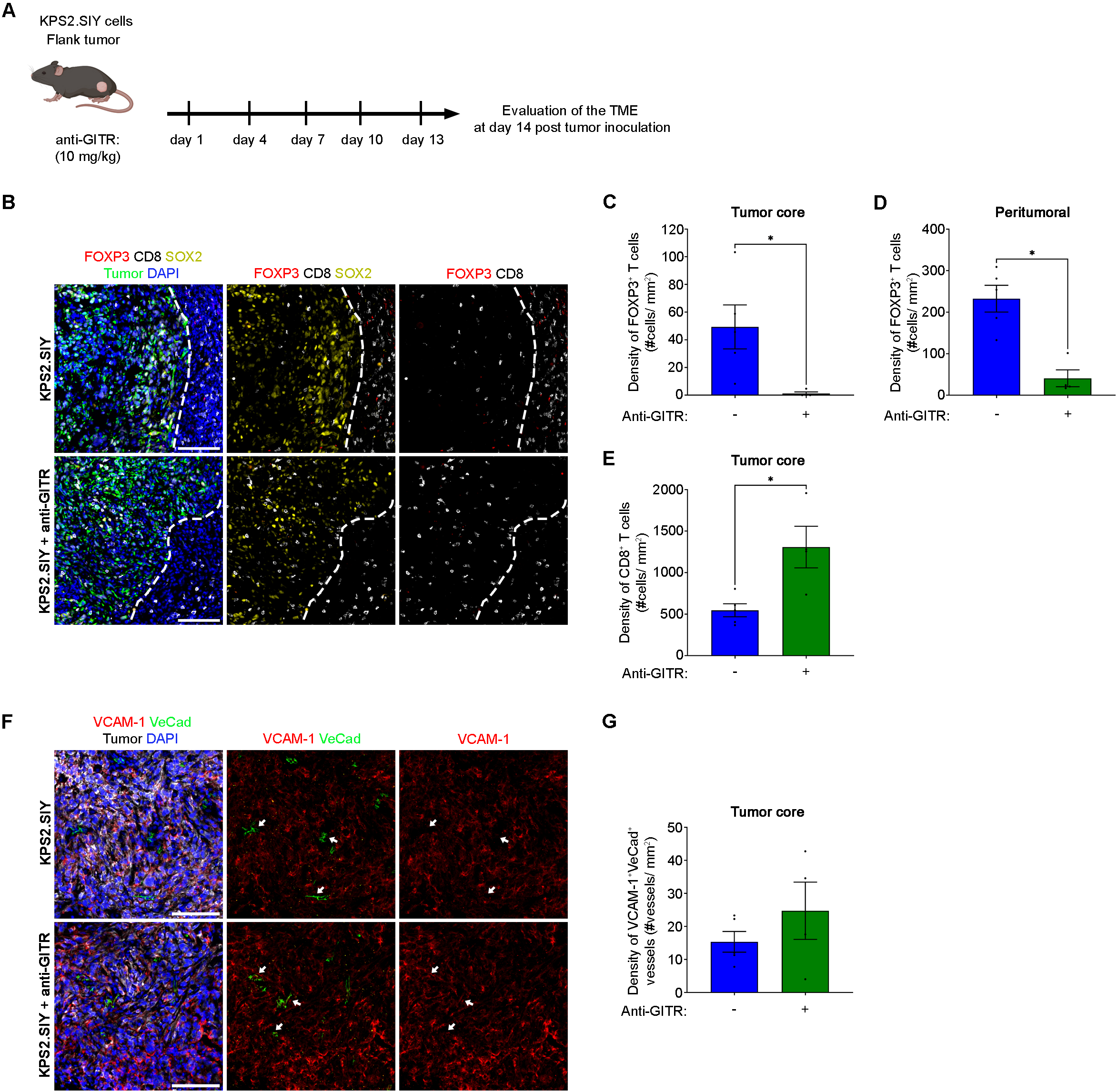
anti-GITR treatment reduces FOXP3^+^ T cells in the TME and improve CD8^+^ T cell infiltration. **A,** Experimental design for (B-G). **B,** Representative fluorescence images of SOX2 (yellow), CD8 (white), FOXP3 (red), and tumor cells (green) in KPS2.SIY and KPS2.SIY+anti-GITR flank tumors at day 14 post-tumor induction. Nuclei were counterstained with DAPI (Blue). Scale bar = 100 μm. **C and D,** Quantification of FOXP3^+^ cell densities in the core and in the peritumoral region of KPS2.SIY and KPS2.SIY+anti-GITR flank tumors. **E,** Quantification of CD8^+^ T cell density in the core of KPS2.SIY and KPS2.SIY+anti-GITR flank tumors. **F,** Representative fluorescence images of VeCAD (green), VCAM-1 (red), and tumor cells (white) in KPS2.SIY and KPS2.SIY+anti-GITR flank tumors at day 14 post-tumor induction. Nuclei were counterstained with DAPI (Blue). Scale bar = 100 μm. **G,** Quantification of VCAM-1^+^VeCAD^+^ vessel density in the core of KPS2.SIY and KPS2.SIY+anti-GITR flank tumors. p-value was determined by Mann Whitney test (N=4-5). Data are shown as means ± SEM. When p-value is significant, results are flagged with stars, *p<0.05.

Our data thus far showed that systemic depletion of Treg cells resulted in restored activation of tumor vasculature (Fig. 5G). We next probed whether anti-GITR mediated changes in the density of Treg cells would similarly affect activation of the tumor vasculature. Strikingly, anti-GITR treatment showed a small yet appreciable increase in the density of activated VCAM-1^+^VeCad^+^ vessels in treated KPS2.SIY tumors compared to untreated KPS2.SIY tumors (24.74 vessels/mm^2^ ± 8.667 SEM treated vs. 15.34 vessels/mm^2^ ± 3.131 SEM untreated; Fig. 6F, 6G). Taken together, our results suggest that reducing Treg cell density via anti-GITR treatment significantly improves CD8^+^ T cell infiltration into the core of KPS2.SIY flank tumors.

### Anti-GITR treatment in combination with checkpoint blockade therapy reduces tumor growth

Our results demonstrate that tumor cell-intrinsic upregulation of SOX2 mediates resistance to CBT due to the exclusion of CD8^+^ T cells from the tumor core. Furthermore, anti-GITR treatment significantly improved CD8^+^ T cell infiltration (Fig. 6B, 6E). Thus, we hypothesized that treating KPS2.SIY flank tumors with anti-GITR in combination with CBT could have synergistic effects and induce tumor regression. To test our hypothesis, we treated KPS2.SIY tumor-bearing mice anti-GITR, CBT, or combination therapy (Fig. 7A). Both single therapies resulted in temporary tumor control, while combination therapy induced complete tumor regression in 4 out of 9 mice (Fig. 7B and Fig. S7A). To exemplify this observation, tumor size at day 21 post-tumor induction was reduced by 69% for both single-agent therapies. In comparison, combination therapy reduced tumor growth by 89% (Fig. 7C), suggesting an additive effect between anti-GITR treatment and CBT.

**Figure 7:**
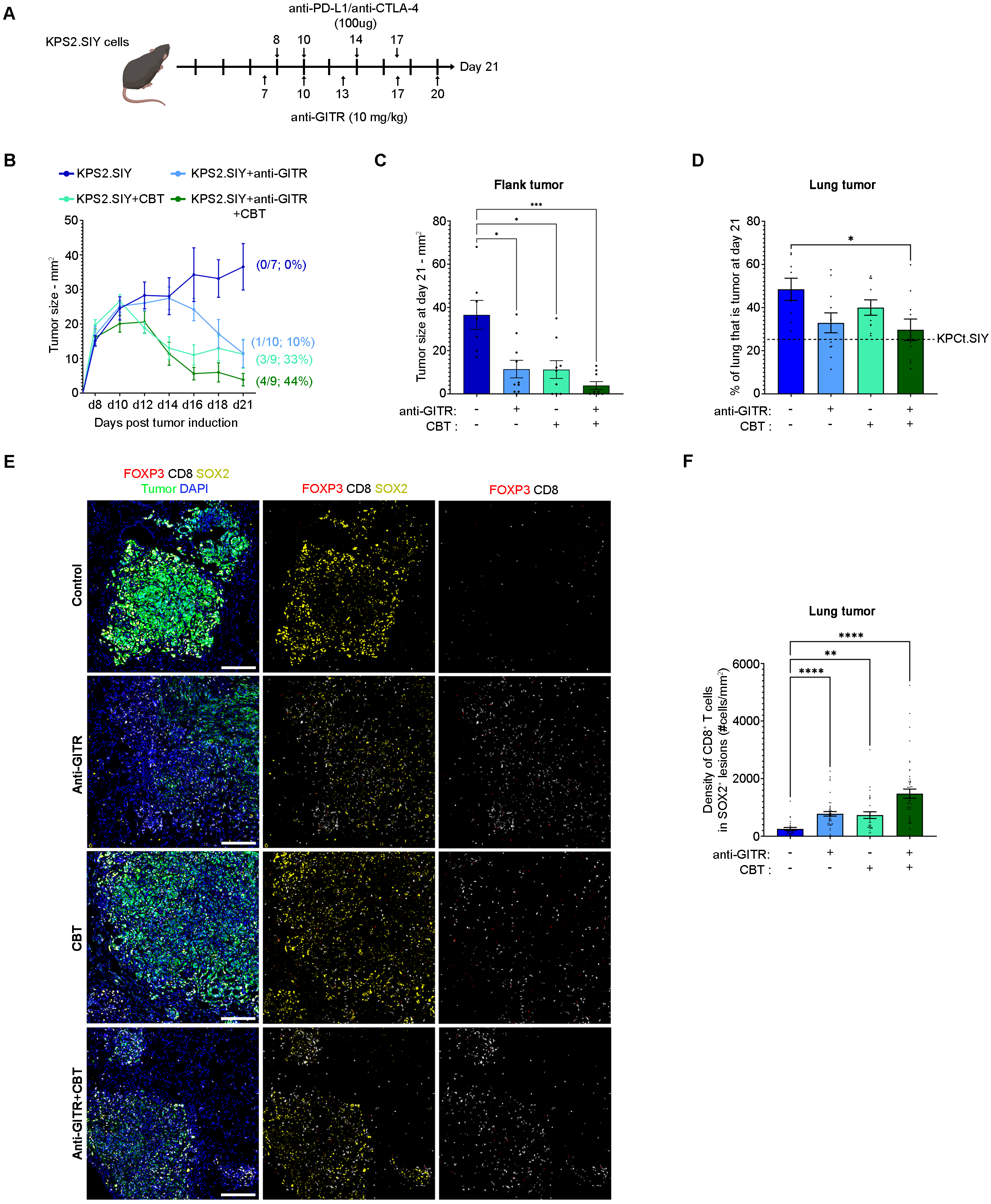
Combining anti-GITR treatment with CBT improves CD8+ T cell infiltration in the tumor and reduces KPS2 flank and lung tumor growth. **A,** Experimental design for (B-F). **B and C,** Flank tumor outgrowth and quantification at day 21 post-tumor induction with the KPS2.SIY cell line and treated with anti-GITR, CBT, anti-GITR+CBT, or control. p-value was determined by Kruskal-Wallis test (N=7-10). **D,** Lung tumor burden at day 21 post inoculation with the KPS2.SIY cell line and treated with anti-GITR, CBT, anti-GITR+CBT, or control. p-value was determined by one-way ANOVA (N=8-11). **E,** Representative fluorescence images of SOX2 (yellow), CD8 (white), FOXP3 (red), and tumor cells (green) in KPS2.SIY lung tumors at day 21 post-tumor induction and treated with anti-GITR, CBT, anti-GITR+CBT, or control. Nuclei were counterstained with DAPI (Blue). Scale bar = 200 μm. **F,** Quantification of CD8^+^ T cell densities in SOX2^+^ lung tumor lesions at day 21 post-tumor induction. p-value was determined by Kruskal-Wallis test (number of lesions=27-46). Data are shown as means ± SEM. When p-value is significant, results are flagged with stars, ****p<0.0001; ***p<0.0005; **p<0.005; *p<0.05.

Because anti-GITR restores CD8^+^ T cell infiltration in flank tumors and combination therapy significantly decreases flank tumor growth, we next assessed the therapeutic effect of the combinatory treatment on orthotopic KPS2.SIY lung tumors. When we assessed the percentage of lung that is tumor after the different therapies, we found that both single treatments did not change tumor burden (Fig. 7D and Fig. S7B). Previous studies by us have shown that KP lung tumors are not responsive to CBT^18^. Surprisingly, we found that when mice were treated with the combination of anti-GITR and CBT, the percentage of the lung that was tumor significantly decreased compared to untreated condition (Fig. 7D and Fig. S7B). To determine changes in CD8^+^ T cell infiltration after single-agent therapies or combination therapy, we analyzed the density of CD8^+^ T cells in SOX2^+^ positive tumor lesions by mIF (Fig. 7E). Consistent with our observations in flank tumors (Fig. 6B, 6E), CD8^+^ T cell density significantly increased after anti-GITR treatment and specifically the combination therapy showed the highest density of CD8^+^ T cells in SOX2^+^ tumor lesions (254.9 cells/mm^2^ ± 50.48 SEM in control vs. 986 cells/mm^2^ ± 132.6 SEM in anti-GITR vs. 627.1 cells/mm^2^ ± 48.93 SEM in CBT vs. 1218 cells/mm^2^ ± 126.3 SEM in anti-GITR+CBT; Fig. 7E, 7F). Furthermore, we also determined Treg cell density in SOX2^+^ lung tumor lesions by mIF after the different therapies. Surprisingly, we found increased Treg cell infiltration after single-agent treatment (Fig. S7C), suggesting a different mechanism of action in flank vs. lung tumors. Taken together, our results show that depletion of Treg cells by anti-GITR treatment restores CD8^+^ T cell infiltration in SOX2^+^ overexpressing flank and lung tumors and combined with CBT significantly reduces tumor burden.

## Discussion

Here, we identified that tumor cell-intrinsic SOX2 expression induces the exclusion of CD8^+^ T cells from the tumor core and promotes resistance to CBT in NSCLC. Mechanistically, tumor cell-intrinsic SOX2 expression increased Treg cell density in the peritumoral region, which prevented activation of the tumor vasculature, thereby blunting infiltration of effector CD8^+^ T cells into the tumor core. We further utilized an anti-GITR therapy to reduce Treg-mediated suppression of the vasculature, resulting in improved CD8^+^ T cell infiltration and response to CBT.

Altered tumor-intrinsic signaling pathways can induce a suppressive TME, facilitating tumor immune evasion and progression^23^. In melanoma, activation of the β-catenin-dependent Wnt signaling pathway in tumor cells blunts recruitment of CD103^+^ DCs to the TME. Consequently, priming and activation of tumor-specific CD8^+^ T cells is reduced, resulting in a lack of T cell infiltration in the tumor^19^. Specifically in NSCLC, the loss of function of *LKB1* is associated with increased infiltration of immunosuppressive neutrophils and reduced T cell infiltration in the TME^8^. In both examples, activation of β-catenin signaling or loss of function of *LKB1* significantly reduces T cell infiltration, resulting in a “cold” type of tumor in both cancer models. Here, we provide evidence of a tumor cell-intrinsic pathway that mediates T cell-exclusion and is associated with poor responses to immunotherapy. Specifically, we showed that high levels of SOX2 in lung cancer cells is associated with T cell exclusion from the tumor core. Previous studies in other cancers have already implied that tumor-intrinsic SOX2 expression suppresses the antitumor immune response through different mechanisms involving tumor cell interferon production by tumor cells^37,38^. While we did not detect any changes in tumor cells to produce interferon, we observed that tumor cell-intrinsic expression of SOX2 affected Treg cells and tumor vasculature resulting in blunted infiltration of CD8^+^ effector T cells. Consistently, Dhodapkar *et al.* detected SOX2-specific CD4^+^ and CD8^+^ T cells in the blood of NSCLC patients with high SOX2 expression^39^, suggesting the capacity of the immune system to induce systemic immune responses against SOX2^+^ tumors. These findings are consistent with our observations that T cell activation is intact in SOX2^+^ tumors. In sum, our results suggest an additional immune evasion mechanism whereby SOX2 promotes resistance to immunotherapy by remodeling the TME and thus preventing T cell infiltration.

The composition of the tumor immune cell compartment plays a crucial role in determining an effective antitumor immune response^40^. Our findings showed a high density of Treg cells in the KPS2 peritumoral region. This Treg cell accumulation was detrimental to CD8^+^ T cell infiltration since Treg cell depletion restored CD8^+^ T cell infiltration and tumor control. Recruitment of Treg cells to the TME by diverse mechanisms has been shown in different cancer types, where they promote a tolerant TME^24^. Interestingly, Xu *et al.* showed that in breast cancer, Treg cells were specifically recruited to the tumor site by CCL1-SOX2-dependent expression in cancer stem cells. As a feedback loop, Treg cells in the TME stimulate the stem cell properties in the tumor, and these changes result in increased tumor progression and metastasis^41^. Crosstalk between Treg cells and stem cell niches have also been shown in steady-stage and regeneration of the skin, the intestine, and the lung, where the presence of Treg cells is crucial for proper tissue repair. Besides, different studies suggest that in cancer, tumor cells recapitulate a regenerative program that allows them to grow^42,43^.

Our data showed that the vasculature in KPS2 flank tumors exhibited characteristics of poor activation. Changes in the tumor vasculature is a hallmark of cancer. Understanding how the tumors induce an aberrant vasculature is crucial to improve T cell infiltration and drug delivery^44^. We observed that KPS2 flank tumors have a reduced expression of VCAM-1, which correlated with the poor CD8^+^ T cell infiltration found in the tumor core. VCAM-1 expression is induced under inflammatory conditions to support leukocyte infiltration into the inflamed tissue^25^. Processes like angiogenesis induce an immune suppressive microenvironment that does not support CD8^+^ T cell extravasation^28^. In this regard, previous data from ovarian cancer showed that Treg cells induce tolerance by promoting tumor angiogenesis^27^. Similarly, our data suggest that Treg cells in KPS2 flank tumors impaired the activation of the endothelial cells, reducing the influx of CD8^+^ T cells into the tumor core. Consistently, our analysis showed that the depletion of Treg cells significantly improved the activation of the tumor vasculature and restored CD8^+^ T cell infiltration. These results are in agreement with previous studies where Treg depletion increased the formation of intratumoral high endothelial venules^45^, which are highly specialized blood vessels that facilitate lymphocyte extravasation^46^. Nevertheless, it remains possible that CD8^+^ T cells are also directly suppressed by Treg cells due to their proximity in the peritumoral region. Further characterization of Treg subpopulations in KPS2 tumors will help elucidate their suppressive role in the TME and understand how these cells modulate the vasculature of SOX2^+^ tumors.

Our data showed that anti-GITR treatment significantly reduces Treg density in the TME, and in combination with anti-PD-L1 and anti-CTLA-4 reduces resistance to CBT in SOX2^+^ tumors. Previous studies demonstrated that Treg cells in the TME express higher GITR levels than Treg cells in the TdLN^34^. Interestingly, clinical and preclinical studies have shown that anti-GITR monotherapy is insufficient to observe significant tumor control^35,47^. Thus, our findings provide a rational for further investigation on the effect of depleting GITR^+^ Treg cells in NSCLC patients with high expression of SOX2 and a CD8^+^ T cell exclusion phenotype, suggesting that combinatory therapy could benefit this particular subset of patients. In summary, our data indicate that in NSCLC, changes in the TME due to dysregulation of tumor-intrinsic SOX2 induce tumor progression by excluding T cells from the tumor core. These changes can be reverted by depleting GITR^+^ Tregs in the TME and in combination with CBT significantly improve tumor control.

## Supporting information

Torres-Mejia Supplement

## Acknowledgments

We thank Rakesh Jain (MGH), Richard Hynes (MIT), and Thomas Diefenbach (Ragon) for experimental guidance. We thank Melissa Duquette for mouse colony maintenance and Paul Thompson for administrative support. We thank the Koch Institute Swanson Biotechnology Center and the Ragon Institute Microscopy Core for technical support. We thank Dr. Walter Newman, Dr. Michael Hass, and Dr. Michael Kagey for experimental guidance and reagents. This work was supported by NCI K99/R00 award, the Ludwig Center at MIT, the SITC-Nektar Therapeutics Equity and Inclusion in Cancer Immunotherapy Fellowship, and Leap Therapeutics Inc.

## Author Contributions

E.T.-M. and SS conceptualized the study. E.T.-M. and SS designed the experiments and interpreted data. E.T.-M., ED, SW, LY performed experiments. E.T.-M., KN, SW analyzed data. E.T.-M. and SS wrote and edited the manuscript. SS and E.T.-M. acquired funding. SS supervised the study.

## Methods

### Mice

Female and male C57BL/6 mice were purchased from Taconic Biosciences or the Jackson Laboratories. Female and male CD45.1^+^ mice were purchased from Taconic Biosciences. Female and male TCR-transgenic 2C CD45.1^+^ and 2C CD45.2^+^ Rag2^-/-^, Rag2^-/-^, and FoxP3^DTR.GFP^ mice were bred and maintained in-house. Experiments were performed using gender-matched 8-14 weeks-old mice. Mice were maintained under specific pathogen-free conditions at the Koch Institute animal facility, and all animal procedures were approved by the Committee on Animal Care (CAC/IACUC) at MIT.

### Generation of expression vectors and modified tumor cell lines

The mouse lung adenocarcinoma cell line KP (Kras^G12D/+^; Trp53^−/−^) was a gift from Prof. Tyler Jacks. The Jacks Laboratory derived this cell line from primary lung tumors from KP C57BL/6 mice^17^. Single-cell clones were generated from the KP cell line by culturing one cell per well in a 96 well/plate. We selected one clonal cell line (KP4C) with low SOX2 protein levels for further experiments. The KP4C clonal cell line was genetically engineered to stably overexpress SOX2 (KPS2 cell line) by retroviral transduction using the pMXs-SOX2-IP construct (Adgene cat#15919). As a control, the pMXs-SOX2-IP construct was linearized by digestion with PacI and EcoRI restriction enzymes (NEB) to cut the SOX2 gene (pMXs-IP). The control cell line KPCt was generated by retroviral transduction of the KP4C clonal cell line using the pMXs-IP construct. After retroviral transduction, KPCt and KPS2 were selected with 20 μg/ml puromycin (GIBCO). The SIYx3-GFP insert was generated with the primers: 5’-GGTGTCGTGAGGATCCACCATGGTGTCTATTTACAGGTAC-3’ 5’-CGCCCTCGAGGAATTCTTACTTGTACAGCTCGTCCATGC-3’ and cloned into pLV-EF1a-IRES-Blast (Addgene #85133) linearized with BamHI and EcoRI restriction enzymes (NEB) using the Clontech In-Fusion HD Cloning System (Clontech #639645). The KPS2 and KPCt were genetically engineered to stably express the model antigen SIYx3-GFP by lentiviral transduction using the pLV-EF1a-SIYx3-GFP-IRES-Blast construct. The newly generated GFP expressing KPS2.SIY and KPCt.SIY cell lines were selected with 2 μg/ml blasticidin (GIBCO). Additionally, both KPCt.SIY and KPS2.SIY cell lines with similar fluorescence intensity levels of GFP were FACS-sorted using FACSAria™ Cell Sorter (BD).

### Tumor cell lines and tumor induction

All cell lines were cultured at 37 °C and 5% CO2 in DMEM (GIBCO) supplemented with 10% heat-inactivated fetal bovine serum, 1X HEPES (GIBCO), and 1%

Penicillin/streptomycin (GIBCO). Cell lines were regularly tested for mycoplasma. For tumor induction, cells were trypsinized with 0.25% of Trypsine and 0.9 mM EDTA for 5 min at 37 °C. Cells were washed three times with PBS and resuspended in PBS (GIBCO). Lung tumors were induced by tail veil injection of 2.5×10^5^ tumor cells in 100 ul of PBS, and flank tumors were generated by subcutaneous injection of 1×10^6^ tumor cells in 100 ul of PBS. Flank tumor growth was quantified by measuring the area of the subcutaneous tumor (calculated by length x width) 3 times a week until the endpoint of the study. Lung tumor burden was determined by analyzing lung tissue sections stained with Hematoxylin and Eosin (H&E) and quantifying the percentage of the lung that was tumor using the QuPath software.

### Virus production

The retroviral packaging cell line Platinum-E was co-transfected with pMXs-SOX2-IP or pMXs-IP using the DNA Transfection Reagent X-TremeGene 9 (ROCHE, Cat# 06365779001) and following the manufacturer’s instructions. The cell line LentiX was co-transfected with the constructs pLV-EF1a-SIYx3-GFP-IRES-Blast, psPAX2 (Addgene#12260), and pDM2.G (Addgene#12259) in Opti-MEM™ I Reduced Serum Medium (ThermoFisher) with Polyethylenimine Linear, MW 25,000 (Polysciences). Virus-containing supernatant was collected at 24 and 48 hours post-transfection and used to transduce tumor cell lines.

### Tissue processing for histology

Flank tumor tissue was collected at different time points after tumor induction and fixed in periodate-lysine-paraformaldehyde buffer (PLP) ^48^ (0.05 M phosphate buffer containing 0.1 M lysine, 2 mg/ml NaIO4, and 1% of paraformaldehyde (Electron Microscopy Grade), pH 7.4) overnight at 4C. Tumor-bearing lungs were perfused with 10 ml of PBS followed by 10 ml of PLP into the heart’s right ventricle. Lungs were dissected and inflated with PLP into the trachea. The tissue was fixed overnight in PLP at 4C. Tissue was cryoprotected by incubation in 15% sucrose in PBS for 24 hours, followed by 30% sucrose at 4C until the tissue sank. Lungs were inflated with 50% optimum cutting temperature compound (OCT) in PBS. Lungs and flank tumors were embedded in 100% OCT and snap-frozen in 2-Methylbutane on dry ice and stored at -80 until sectioning. Frozen tissue was sliced at 10 μm using a Cryostar NX70 (Thermo Scientific). Frozen tissue sections were dried at room temperature for 1 hour and post-fixed in ice-cold acetone for 10 min at -20C; samples were dried for 1 hour and stored at -20C until processing. Flank and lung tumors were dissected and fixed in 10% formalin (Sigma) for 72 hours at room temperature for paraffin-embedded tissue. Samples were transferred to 70% ethanol for 24 hours and then dehydrated by incubation in 95% ethanol, 100% ethanol, and Xylene. Tissue was embedded in paraffin and sliced at 5 μm.

### Tissue staining

Paraffin-embedded sections were first warmed in a slide warmer at 55C for 20 minutes. Samples were dewaxed by incubating the sections in Xylene for 3 minutes, 3 times, then 3 minutes in 100% ethanol, 1 minute in 95% ethanol, 1 minute 70% ethanol, and in running water, 1 minute twice. For hematoxylin and eosin staining, tissue sections were treated as follows: 3 minutes in Harris Acidified Hematoxylin, 1 minute in running water, 5 seconds acid alcohol, 1 min running water, 1 minute bluing, 1 minute in running water, 1 minute in 95% ethanol, 10 seconds in Eosin, 1 minute in 95% ethanol, 1 minute (twice) in 100% ethanol, 2 min (three times) in Xylene, and mounted using Tissue-Tek® Glas™ Mounting Medium (Sakura). For immunofluorescence staining, sections were subjected to antigen retrieval by incubation in citrate buffer using the 2100 Retriever following the manufacturer’s instructions (Electron Microscopy Science). Samples were incubated for 2 hours at room temperature using the blocking solution: Blocker Casein in tris-buffered saline (ThermoScientific, Cat# 37532), 0.3% Triton X-100, and 10% normal donkey or goat serum (Jackson Immunoresearch). Samples were incubated with primary antibodies diluted in blocking solution overnight at 4C in a moister chamber. Samples were washed with PBS and 0.1% Triton X-100 and incubated for 1 hour at room temperature with secondary conjugated antibodies diluted in the blocking solution. Samples were washed 3 times with PBS and incubated for 30 seconds with TrueBlack following the manufacturer’s instructions (Biotium, Cat# 23007). Samples were washed with PBS, and Nuclei were counterstained using DAPI (Sigma). Finally, the samples were mounted using ProLong Diamond Antifade Mountant (Thermo Fisher Scientific, Cat#P36970) and curated for 24 hours before imaging.

Frozen sections were warmed at room temperature for 30 minutes and then washed with PBS. Samples were blocked and permeabilized in PBS containing 1% bovine serum albumin (Sigma), 2% normal mouse serum (Sigma), 10% normal donkey or goat serum, and 0.3% Triton X-100 (blocking solution for frozen sections) for 2 hours at room temperature. Tissue was incubated with primary antibodies diluted in blocking solution for frozen sections overnight at 4C in a moister chamber. Samples were washed with 0.1% Triton X-100 in PBS three times and incubated with secondary antibodies diluted 1:500 in blocking solution for frozen sections for 1 hour at room temperature in a dark chamber. Samples were washed with 0.1% Triton X-100 in PBS three times. Sections stained with fluorophore-conjugated primary antibodies were incubated with the fluorophore-conjugated antibody diluted in the blocking solution for frozen sections for 1 hour at room temperature in a dark chamber. Nuclei were counterstained using DAPI (Sigma) or TO-PRO-3 (ThermoFisher). Samples were mounted using ProLong Diamond Antifade Mountant and curated for 24 hours before imaging. A list of antibodies is provided in Table 2.

### Multiplex immunofluorescence imaging (mIF) and quantitative analyses

Tissue sections were imaged using the TissueFAXS Plus automated slide scanning system (TissueGnostics GmbH, Austria) combining a Zeiss Axio Imager 2 upright microscope with a Märzhäuser motorized stage (Märzhäuser Wetzlar). Images were acquired using a Zeiss 20x Plan-Neofluor 0.5 NA air objective with the following filters sets AF488 (470/24) ET470/30x T495lpxr ET515/30m, AF750 (740/20) ET740/40x T770lpxr ET780lp, multiband dichroic qTexasRed/qCy5 (550/15, 640/30) from Chroma Technology, USA. Excitation was provided by a Lumencor Spectra 3 LED light engine (500 mW per channel). Fluorescence images were captured using a Hamamatsu Orca Flash 4.0 V2 cooled digital CMOS camera C11440-22CU. Tissue sections were acquired as z-stacks in a process called extended focus, with a 2.5 μm step size, including one step above and one step below the focal plane. Extended focus takes each in-focus area of each image within the z-stack and combines those regions into a single image. Image processing, including stitching, was performed using the TissueFAXS capture/control software.

Cell densities were quantified using TissueQuest image analysis software (TissueGnostics USA). Briefly, regions of interest (ROI) were defined in the core and in the peritumoral area. Automatic segmentation was performed in TissueQuest based on DAPI nuclear staining. The density of positive cells was calculated in the different ROI by the number of positive events divided by the ROI area. Lung tumor burden and vessel density were analyzed using the QuPath software for digital pathology image analysis^49^. We first created a threshold for lung tumor burden to detect the long lobe to analyze. We then defined tumor and non-tumor cells based on the lung H&E staining. Finally, the percentage of the lung that was tumor was calculated as the number of tumor cells divided by the total number of events counted in one lung lobe. Vessel density was determined based on VeCad staining in the core of flank tumors (Fig. S6). Tumor regions in the flank tissue sections were defined based on the expression of GFP by the tumor cell lines or panKeratin positive staining. Vessel detection was performed by creating a threshold to detect VeCad-positive staining in the tumor core. Them, we classified the vessels depending on the expression of VCAM-1 as positive or negative. Vessel density was determined as the number of vessels divided by the defined tumor region. Image postprocessing was performed with the image processing package FIJI^50^.

### Heat map

Gene expression data set (release date February 4, 2015) for NSCLC (LUAC and LUSC) was downloaded from TCGA and processed as described by Spranger et al.^14^. We used a list of 160 genes for clustering NSCLC (LUAD and LUSC) patients in high, medium or low T cell expression^14^. Expression data were log-transformed via a log2(x+1) transformation. The heatmap.2 function from the “gplots” (version 3.1.1) R package was used to perform hierarchical clustering of the genes and patient tumors and visualization of the clusters using Euclidean distance metrics and Ward agglomerative clustering (using option “ward.D2”). Program: RStudio Version 1.4.1106.

### Unbiased pathway analysis

We first cluster patients into T cell infiltrated and non-T cell infiltrated groups. Then, we determined the differentially expressed genes between both groups (cluster 1-3 T cell infiltrated and cluster 6-8 non-T cell infiltrated) as previously described^19^. Initial pathway analysis was performed using the Ingenuity Platform, which identified SOX2 as a potential candidate pathway. SOX2 expression levels were subsequently assessed per patient using normalized gene expression values.

### Tissue dissociation

To analyze immune cells in the lung TME, we performed retro-orbital injections with an anti-CD45 fluorescence-labeled antibody to discriminate circulating immune cells. Mice were sacrificed 5 min post injections by cervical dislocation, and lung tissue was collected for further processing. Flank and lung tissues were dissected, minced, and enzymatically dissociated by incubation in 250 μg/ml Liberase (Sigma-Aldrich) and 50 μg/ml DNase (Sigma-Aldrich) in RPMI (GIBCO) for 45 minutes at 37C. Tissue was mashed through a 70 μm filter and washed with PBS. Dissociated lung samples were layered over Ficoll (GE) and spun at 450g for 30 minutes with cero acceleration and brake. Immune cells located in the interphase of PBS and Ficoll were collected and washed with PBS. Lung and flank dissociated cells were washed with chilled FACS buffer (PBS, 2% FBS, Brefeldin A, and 2 mM EDTA) and kept at 4C for further analysis. Spleens and TdLN were directly mashed through a 70 μm filter into RPMI followed by a PBS wash. Spleens were incubated in ACK buffer for 3 minutes on ice and then washed with PBS. To analyze tumor cells in the lung or in the flank, tissue was minced and enzymatically dissociated by incubation in DMEM/F12, HEPES, DNAse, Collagenase from *Clostridium histolyticum* (167μg/ml), CaCl2 (0.06mM) for 30 minutes at 37C. Tissue was mashed through a 100 μm filter and washed with PBS. Dissociated cells were incubated with ACK buffer for 3 minutes at room temperature and washed with PBS. Tumor cells were resuspended in FACS buffer and kept at 4C for further analysis.

### Flow cytometry

Cells were incubated with anti-CD16/CD32 for FC block diluted in FACS buffer for 15 minutes at 4C. Cells were washed with FACS buffer and stained for surface proteins using fluorophore-dye conjugated antibodies diluted in FACS buffer for 20 min at 4C (Antibody list in Table S2) together with fixable live-dead staining. Cells were washed with FACS buffer followed and directly analyzed by flow cytometry or fixed for intracellular staining. The Foxp3 Transcription Factor Fixation/Permeabilization buffer (eBioscience) was used following the manufacturer’s instructions for intracellular staining. Brefeldin A (BioLegend) was added to buffers before fixation for cytokine staining. After fixation, cells were washed three times with FACS buffer and stained for intracellular proteins in FACS buffer overnight at 4°C. Cell suspension was washed twice with FACS buffer prior to flow analysis. Precision Count Beads (BioLegend) were added to samples following the manufacturer’s instructions to obtain absolute counts of cells. Flow cytometry sample acquisition was performed on an LSR Fortessa cytometer (BD). Flow cytometry data were analyzed using FlowJo software (TreeStar). For cell sorting, staining was performed as described above under sterile conditions, and samples were sorted using a FACSAria™ Cell Sorter (BD).

### IFNγ-ELISpot

ELISpot plates (EMD Millipore) were coated overnight at 4°C with anti-IFNγ capture antibody (BD Biosciences) diluted 1:200 in PBS. Plates were washed and blocked with DMEM (GIBCO) supplemented with 10% FBS (Atlanta Biologicals) and 1% penicillin/streptomycin (GIBCO) for 2 hours at room temperature. Splenocytes isolated as described above and seeded at a density of 1×10^6^ cells/well in the presence or absence of 160 nM SIY peptide in DMEM, supplemented with 10% FBS, 1% penicillin/streptomycin. As a positive control, splenocytes were incubated with 100 ng/mL Phorbol 12-myristate 13-acetate (Sigma-Aldrich) and 1 μg/mL ionomycin (Sigma-Aldrich). Cells were incubated overnight at 37°C and 5% CO2. Developing was performed using the IFNγ-ELISpot kit (BD Biosciences) and following the manufacturer’s instructions. Plates were imaged and analyzed using the CTL ELISPOT -ImmunoSpot®.

### In vivo proliferation assay

*In vivo* proliferation assay was adapted from Horton et al. 2021^18^. Briefly, splenocytes were isolated from the spleen of TCR-transgenic 2C CD45.1^+^ or 2C CD45.2^+^ Rag2^-/-^ mice, as described above. Splenocytes were labeled with CellTrace™ CFSE Cell Proliferation Kit following the manufacturer’s instructions (Invitrogen). 1×10^6^ of labeled splenocytes were adoptively transferred by retro-orbital injection of a flank or lung tumor-bearing mice 7 days post-tumor induction. Recipient mice were euthanized, and TdLN and spleen were collected 3 days after SIY-reactive CD8^+^ T cells adoptive transfer. Tissue was dissociated, and the proliferation of transferred SIY-reactive CD8^+^ T cells was analyzed by flow cytometry. The percentage of proliferated SIY-reactive CD8^+^ T cells was determined by the dilution of CFSE stain due to rounds of proliferation. As a negative control of proliferation (maximum signal of CFSE), labeled splenocytes were kept in culture for 72 without *in vitro* activation and used to gate on the undivided cell population.

### Ex-vivo activation of SIY-reactive T cells

SIY-reactive CD8^+^ T cells were isolated from the spleen and lymph nodes from TCR-transgenic 2C CD45.1^+^ or 2C CD45.2^+^ Rag2^-/-^ mice using the CD8a^+^ T cell isolation kit (Miltenyi Biotec), following the manufacturer’s instructions. Isolated CD8^+^ T cells were washed three times with PBS and seeded at a density of 5×10^5^ cells/cm^2^ in a plated pre-coated with 0.2 μg/ml anti-CD3 and 0.5 μg/ml anti-CD28. Isolated CD8^+^ T cells were cultured for 72 hours in T cell culture media: RPMI (GIBCO, Cat#11875-093) supplemented with 10% of heat-inactivated FBS, 1% penicillin/streptomycin (GIBCO, Cat#15140-122), and 1:1000 2-Mercaptoetanol (Gibco, Cat#21985-023).

### Effector SIY-reactive T cell adoptive transfer

Flank tumors were generated by subcutaneous injection of 1×10^6^ KPCt.SIY or KPS2.SIY cell lines. *Ex-vivo* activated 1×10^6^ SIY-reactive CD8^+^ T cells were injected retro-orbitally to a flank tumor-bearing recipient mouse, 7 days post-tumor induction. Flank tumor tissue was collected 3 days after adoptive transfer and fixed in PLP buffer overnight at 4°C.

### T cell in vitro killing assay

KPCt.SIY and KPS2.SIY were co-cultured with non-SIY expressing parental cells KPCt and KPS2, respectively. Parental cells were previously labeled with Cell Trace Violet Proliferation kit, following the manufacturer’s protocol (ThermoFisher Scientific, Cat# C34557). Cells were seeded in a 24 well/plate at a density of 200.000 cells per well (100.000 cells parental plus 100.000 cells SIY expressing cells) in T cell culture media. After 7 hours post-seeding, *ex-vivo* activated SIY-reactive T cells were added to the tumor cells at different ratios. The percentage of killing was analyzed by flow cytometry 17 hours post-co-culture of tumor cells with SIY-reactive T cells. For flow cytometry analysis, samples were labeled with anti-CD8 antibody and fixable live dead dye APC-Cy7. Single-cell line and SIY-reactive T cell culture were used as control and for gating strategies. The percentage of live cells was calculated as the percentage of live cells normalized to control = (%GFP cells/%Violet Tracer cells) *100 / control sample.

### In vivo mouse treatments

For checkpoint blockade therapy, tumor-bearing mice were injected intraperitoneally with 100 μg of anti-CTLA-4 (clone UC10-4F10-11, Bio X Cell) and 100 μg of anti-PD-L1 (clone 10F.9G2, Bio X Cell) antibodies on days 8, 10, 14, and 17 post-tumor induction. For FoxP3 cell depletion, *FoxP3^DTR.GFP^* mice were injected intraperitoneally, with 1 μg of diphtheria toxin (Sigma-Aldrich) diluted in PBS, on days 7 and 10 post flank tumor induction. Anti-GITR (mDTA-1) treatment was administered intraperitoneally at 10 mg/kg starting on days 1, 4, 7, 10, and 13 post-tumor induction. When anti-GITR was administered in combination with checkpoint blockade therapy, anti-GITR was administered on days 7, 10, 13, 17, and 20 post-tumor induction.

### Statistical analysis

Statistical analysis was performed using the software GraphPad Prism (GraphPad). Nonparametric tests were used for small sample sizes ≤ 4 and for data that did not follow a normal distribution. Normal distribution was determined by a Shapiro-Wilk test and Kolmogorov-Smirnov test. Statistical analyses were performed by Mann-Whitney U or t-student test for comparison of two groups. A Kruskal-Wallis test with a Dunn’s posthoc test, or a one-way ANOVA for multiple comparisons with a Tukey HSD (Honestly Significant Difference) posthoc test were performed for more than two groups. We used two-way ANOVA to determine how two or more variables affected our group of studies. The correlation of CD8^+^ T infiltration and SOX2 expression in NSCLC tissue biopsies were performed by a Fisher’s exact test. Unless otherwise indicated, data are shown as means ± SEM, and differences with a p-value < 0.05 were considered significant.

## Supplementary figure legends

Fig. S1: LUSC has higher expression of SOX2 and low density of CD8^+^ TIL compared to LUAD.

**A,** Representative H&E and fluorescence images of a biopsy showing one selected tumor region and its segmentation for SOX2^+^ cells and CD8^+^ T cells using the TissueQuest Analysis Software (TissueGnostic); scale bar = 2000 μm. **B,** Violin plot shows SOX2 MFI intensity in tumor regions of LUSC and LUAC biopsies. **C,** Violin plot shows CD8^+^ TIL density in selected LUSC and LUAC tumor regions. p-value was determined by Mann-Whitney test. When p-value is significant, results are flagged with stars, ****p<0.0001; **p<0.005.

Fig. S2: Frequency of CD8^+^, CD4^+^, and FOXP3^+^ T cells in whole tumor tissue is similar independently of SOX2 expression in tumor cells.

**A,** Diagram shows the experimental design to generate the lung tumor cell lines: KP4C, KPCt, KPS2, KPCt^Low^, KPS2^High^. **B,** Representative flow plots of SOX2 expression in KP parental, KPCt, KPS1, KPCt^Low^, and KPS2^High^ cell lines. **C and D,** Outgrowth of KPCt^Low^ and KPS2^High^ flank tumors in WT (C56BL/6) and Rag2^-/-^ mice. p-value at day 21 was determined by Mann-Whitney test (N=6). **E,** Frequency of CD8^+^, CD4^+^, and FOXP3^+^ T cells in flank tumors at day 21 post-tumor induction. p-value was determined by t-student (N=9). **F,** Representative flow plot gating strategy for CD8^+^, CD4^+^, and FOXP3^+^ T cells. **G,** Quantification of total number of CD8^+^, CD4^+^, and FOXP3^+^ T cells in the flank tumor tissue. p-value was determined by t-student (N=9). **H,** Quantification and representative flow plots of PD-1 expression in CD8^+^, CD4^+^, and FOXP3^+^ T cells in flank tumors at day 21 post-tumor implantation. p-value was determined by t-student (N=9). Data are shown as means ± SEM. When p-value is significant, results are flagged with stars, ***p<0.0001; *p<0.05.

Fig. S3: Innate immune cells equally infiltrate KPCt and KPS2 tumors.

**A and B,** Representative flow plots and quantification of frequency (A) and PD-1 expression (B) in CD8^+^, CD4^+^, and FOXP3^+^ T cells from lung tumors at day 21 post-tumor implantation. p-value was determined by t-student (N=12-13). **C,** Representative fluorescence images of SOX2 (red), CD45 (White), and panKeratin (green) in KPCt and KPS2 lung tumors at day 21. Nuclei were counterstained with DAPI (Blue). Scale bar = 100 μm. **D,** Quantification of CD45^+^ cell density in lung tumor bearing mice. p-value was determined by t-student (N=5-6). **E,** Representative fluorescence images of SOX2 (red), CD11c (White), and panKeratin (green) in KPCt and KPS2 lung tumors at day 21. Nuclei were counterstained with DAPI (Blue). Scale bar = 100 μm. **F,** Quantification of CD11c^+^ cell density in lung tumor bearing mice. p-value was determined by t-student (N=5). Data are shown as means ± SEM. When p-value is significant, results are flagged with stars, **p<0.005; *p<0.05.

Fig. S4: Priming and activation of CD8^+^ T cells is not affected by overexpression of SOX2 in tumor cells.

**A,** Representative flow plots of SOX2 and SIY-GFP expression in KPCt.SIY and KPS2.SIY cell lines. **B to D,** Representative flow plots and frequency of (B) endogenous SIY-reactive T cells, (C) PD-1^+^ SIY-reactive T cells, and (I) GzmB^+^ SIY-reactive T cells in KPCt.SIY and KPS2.SIY flank tumors at day 8 post-tumor induction. p-value was determined by t-student (N=7). **E and F,** Representative flow plots and frequency of endogenous SIY-reactive T cells and (F) PD-1^+^ SIY-reactive T cells in KPCt.SIY and KPS2.SIY lung tumors at day 8 post-tumor induction. p-value was determined by t-student (N=12). **G,** Experimental design for Fig. 4F, 4G and Fig. S4H. **H,** Representative flow plots of CFSE-labeled 2C T cells and their PD-1 expression in the mTdLN of KPCt.SIY and KPS2.SIY lung-tumor bearing mice at 72 hours after adoptive transfer, day 10 of tumor growth. **I and J,** IFNγ-ELISpot quantification of splenocytes from flank (I) and lung (J) tumor-bearing mice at day 8 post-tumor implantation. p-value was determined by Mann-Whitney test (flank N=5, lung N=4). Data are shown as means ± SEM. When p-value is significant, results are flagged with stars, ***p<0.0005.

Fig. S5: Treg cells are recruited to the peritumoral region of KPS2.SIY flank tumors.

**A,** Representative fluorescence images of SOX2 (cyan), FOXP3 (magenta), and tumor cells (green) in KPCt.SIY and KPS2.SIY flank tumors at day 10 post-tumor induction. Nuclei were counterstained with DAPI (Blue). Scale bar = 100 μm. **B to E,** Quantification of FOXP3^+^ T cell and CD8^+^ T cell density in the tumor core and in the peritumoral region of KPCt.SIY and KPS2.SIY flank tumors. p-value was determined by Mann-Whitney test (N=13). Data are shown as means ± SEM. When p-value is significant, results are flagged with stars, **p<0.005.

Fig. S6: VCAM-1 expression is lower in the TME of KPS2.SIY compared to KPCt.SIY flank tumors.

**A,** KPCt.SIY, KPS2.SIY and KPS2.SIY+DT flank tumors at day 14 post-tumor induction. p-value was determined by one-way ANOVA (N=6-8). **B,** Representative flow plots and quantification of Treg depletion after DT treatment, graph show the frequency of Treg cells in the spleen. p-value was determined by Mann-Whitney test (N=6). **C,** Representative images of the TME of KPCt.SIY, KPS2.SIY, and KPS2.SIY+DT flank tumors at day 14 post-tumor induction, showing mIF for SOX2, CD8, FOXP3, and the tumor, scale bar = 300 μm. **D,** Quantification of SOX2^+^ tumor cell density in KPCt.SIY, KPS2.SIY and KPS2.SIY+DT flank tumors. p-value was determined by one-way ANOVA (N=3-8). **E,** Representative flow plot and histogram showing VCAM-1 expression in KPCt.SIY and KPS2.SIY tumor cells culture in vitro. **F,** example of vessel segmentation analysis based on VeCad expression and classification of vessels depending on VCAM-1 expression. Data are shown as means ± SEM. When p-value is significant, results are flagged with stars, ****p<0.0001; **p<0.005; *p<0.05.

Fig. S7: Anti-GITR treatment makes SOX2^+^ tumors responsive to CBT.

**A,** Flank tumor outgrowth curves generated with the KPS2.SIY tumor cell line and treated with anti-GITR, CBT, anti-GITR+CBT, or control. p-value was determined by two-ways ANOVA (N=7-10). **B,** Representative H&E showing lung tumors generated with the KPS2.SIY cell line and treated with anti-GITR, CBT, anti-GITR+CBT, or control. Scale bar = 1500 μm. **C,** Quantification of FOXP3^+^ T cell densities in SOX2^+^ lung tumor lesions at day 21 post-tumor induction. p-value was determined by Kruskal-Wallis test (Number of lesions=13-37). Data are shown as means ± SEM. When p-value is significant, results are flagged with stars, *p<0.05; **p<0.01.

**Table S1:** NSCLC patient characteristics.

| **Variable** | **LUSC** | **LUAC** |
| --- | --- | --- |
| Number of samples | 12 | 27 |
| Gender (n; %) |  |  |
| Female | 6; 50% | 16; 59.2% |
| Male | 6; 50% | 11; 40.7% |
| Stage (n; %) |  |  |
| I&II | 8; 66.6% | 18; 66.6% |
| III&IV | 4; 33.3% | 9; 33.3% |
| SOX2 positive (n; %) | 11; 91.6% | 9; 33.3% |

**Table S2:**
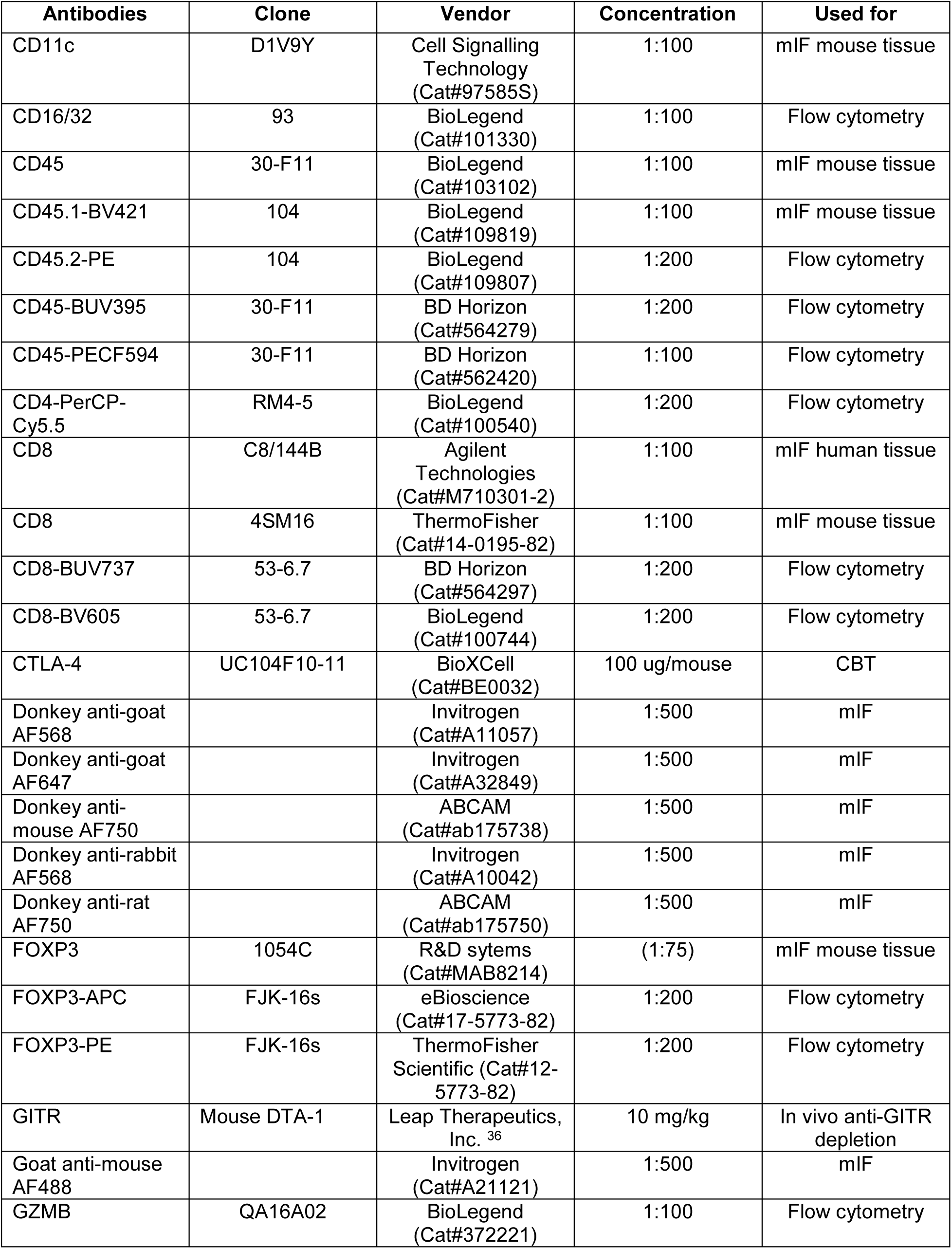
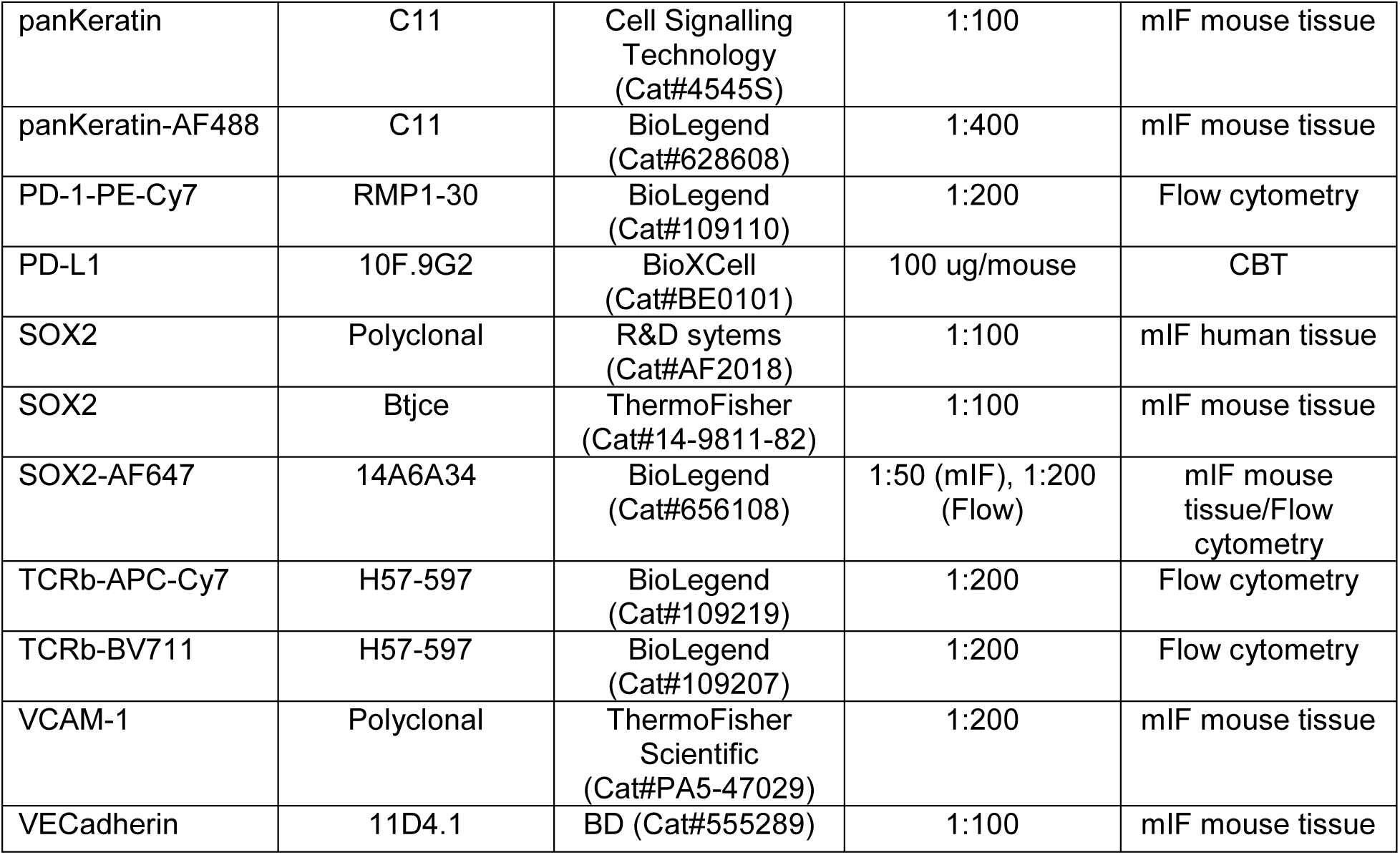
Antibody list.

| **Antibodies** | **Clone** | **Vendor** | **Concentration** | **Used for** |
| --- | --- | --- | --- | --- |
| CD11c | D1V9Y | Cell Signalling Technology (Cat#97585S) | 1:100 | mIF mouse tissue |
| CD16/32 | 93 | BioLegend (Cat#101330) | 1:100 | Flow cytometry |
| CD45 | 30-F11 | BioLegend (Cat#103102) | 1:100 | mIF mouse tissue |
| CD45.1-BV421 | 104 | BioLegend (Cat#109819) | 1:100 | mIF mouse tissue |
| CD45.2-PE | 104 | BioLegend (Cat#109807) | 1:200 | Flow cytometry |
| CD45-BUV395 | 30-F11 | BD Horizon (Cat#564279) | 1:200 | Flow cytometry |
| CD45-PECF594 | 30-F11 | BD Horizon (Cat#562420) | 1:100 | Flow cytometry |
| CD4-PerCP-Cy5.5 | RM4-5 | BioLegend (Cat#100540) | 1:200 | Flow cytometry |
| CD8 | C8/144B | Agilent Technologies (Cat#M710301-2) | 1:100 | mIF human tissue |
| CD8 | 4SM16 | ThermoFisher (Cat#14-0195-82) | 1:100 | mIF mouse tissue |
| CD8-BUV737 | 53-6.7 | BD Horizon (Cat#564297) | 1:200 | Flow cytometry |
| CD8-BV605 | 53-6.7 | BioLegend (Cat#100744) | 1:200 | Flow cytometry |
| CTLA-4 | UC104F10-11 | BioXCell (Cat#BE0032) | 100 ug/mouse | CBT |
| Donkey anti-goat AF568 |  | Invitrogen (Cat#A11057) | 1:500 | mIF |
| Donkey anti-goat AF647 |  | Invitrogen (Cat#A32849) | 1:500 | mIF |
| Donkey anti-mouse AF750 |  | ABCAM (Cat#ab175738) | 1:500 | mIF |
| Donkey anti-rabbit AF568 |  | Invitrogen (Cat#A10042) | 1:500 | mIF |
| Donkey anti-rat AF750 |  | ABCAM (Cat#ab175750) | 1:500 | mIF |
| FOXP3 | 1054C | R&D sytems (Cat#MAB8214) | (1:75) | mIF mouse tissue |
| FOXP3-APC | FJK-16s | eBioscience (Cat#17-5773-82) | 1:200 | Flow cytometry |
| FOXP3-PE | FJK-16s | ThermoFisher Scientific (Cat#12-5773-82) | 1:200 | Flow cytometry |
| GITR | Mouse DTA-1 | Leap Therapeutics, Inc. <sup>36</sup> | 10 mg/kg | In vivo anti-GITR depletion |
| Goat anti-mouse AF488 |  | Invitrogen (Cat#A21121) | 1:500 | mIF |
| GZMB | QA16A02 | BioLegend (Cat#372221) | 1:100 | Flow cytometry |
| panKeratin | C11 | Cell Signalling Technology (Cat#4545S) | 1:100 | mIF mouse tissue |
| panKeratin-AF488 | C11 | BioLegend (Cat#628608) | 1:400 | mIF mouse tissue |
| PD-1-PE-Cy7 | RMP1-30 | BioLegend (Cat#109110) | 1:200 | Flow cytometry |
| PD-L1 | 10F.9G2 | BioXCell (Cat#BE0101) | 100 ug/mouse | CBT |
| SOX2 | Polyclonal | R&D sytems (Cat#AF2018) | 1:100 | mIF human tissue |
| SOX2 | Btjce | ThermoFisher (Cat#14-9811-82) | 1:100 | mIF mouse tissue |
| SOX2-AF647 | 14A6A34 | BioLegend (Cat#656108) | 1:50 (mIF), 1:200 (Flow) | mIF mouse tissue/Flow cytometry |
| TCRb-APC-Cy7 | H57-597 | BioLegend (Cat#109219) | 1:200 | Flow cytometry |
| TCRb-BV711 | H57-597 | BioLegend (Cat#109207) | 1:200 | Flow cytometry |
| VCAM-1 | Polyclonal | ThermoFisher Scientific (Cat#PA5-47029) | 1:200 | mIF mouse tissue |
| VECadherin | 11D4.1 | BD (Cat#555289) | 1:100 | mIF mouse tissue |

