## Supplementary material for "Lung cancer-intrinsic SOX2 expression mediates resistance to checkpoint blockade therapy by inducing Treg-dependent CD8^+^ T cell exclusion": Torres-Mejia Supplement

### SUPPLEMENTARY FIGURE 1

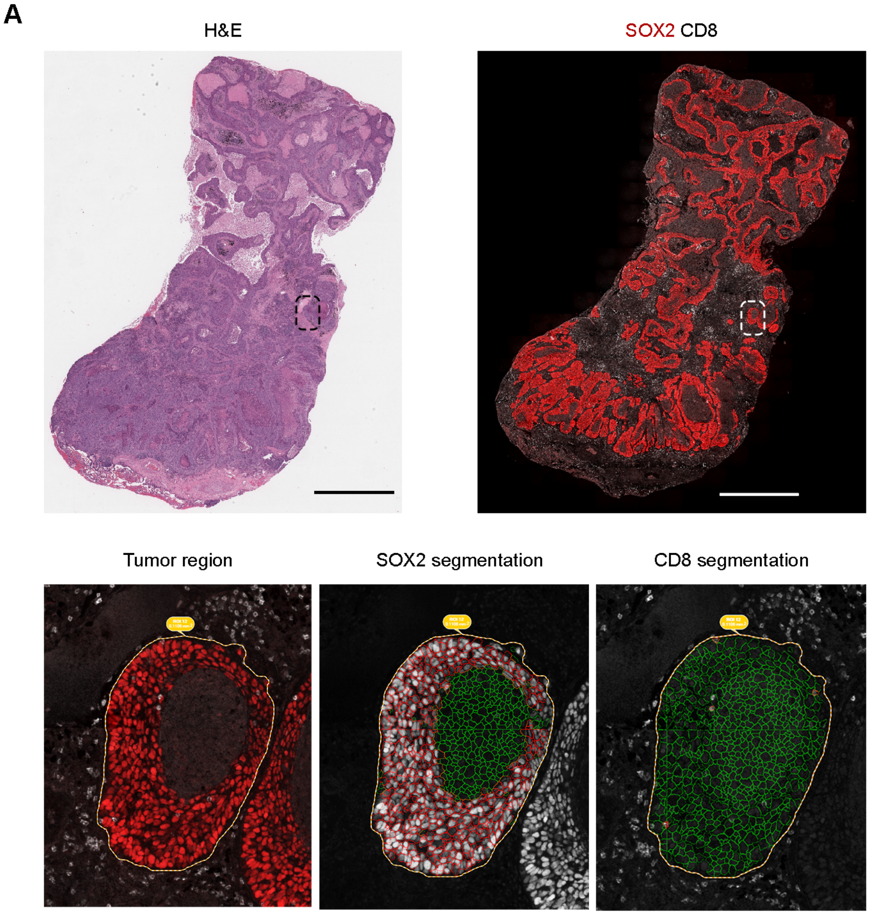


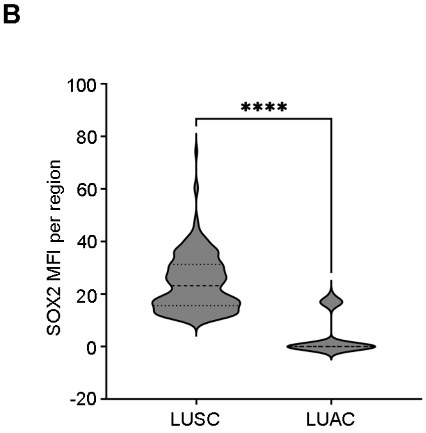


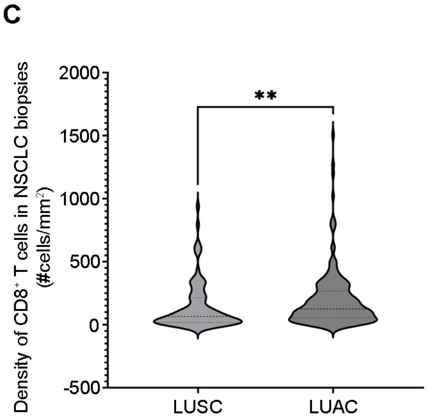

SUPPLEMENTARY FIGURE 2

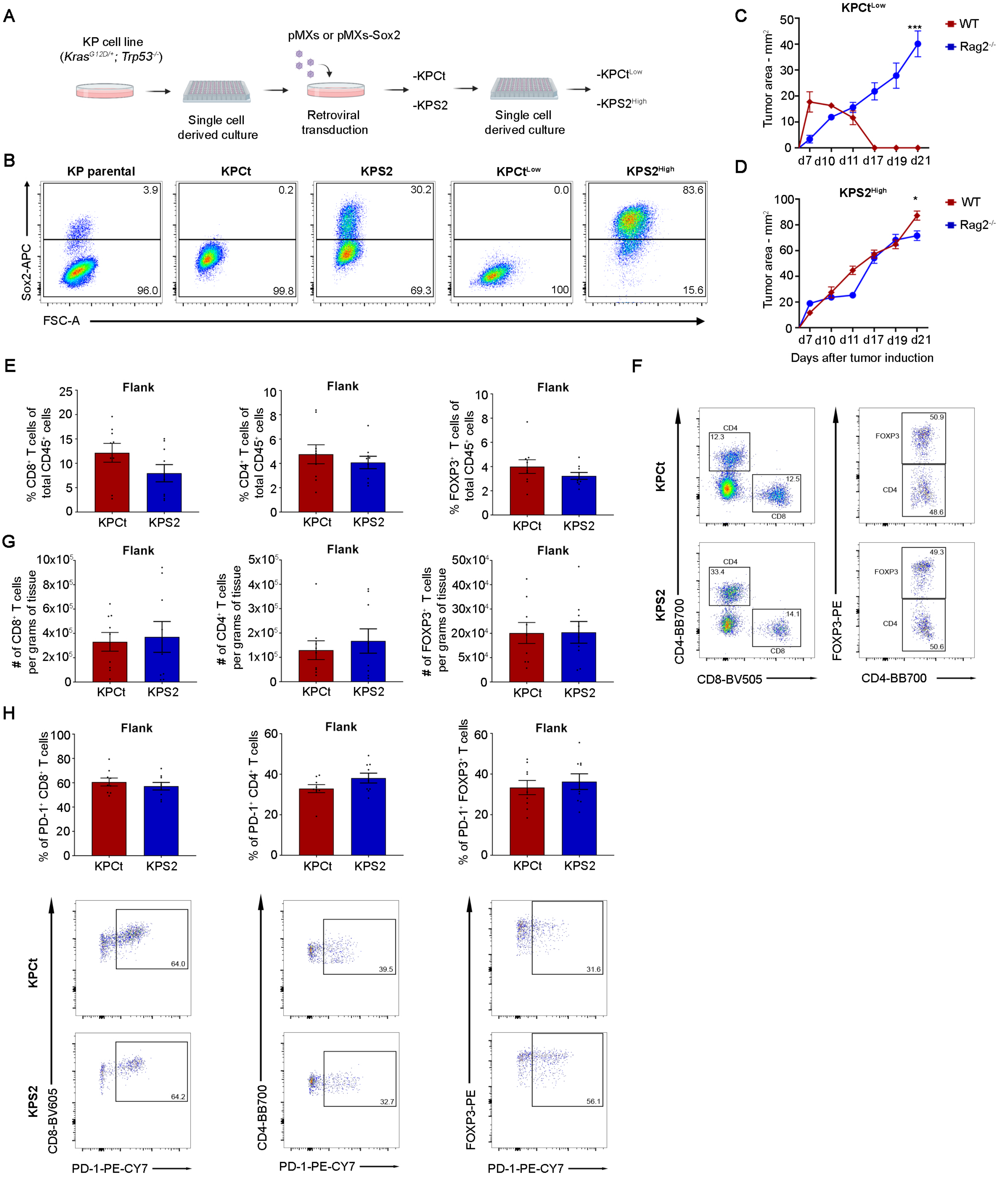

### SUPPLEMENTARY FIGURE 3

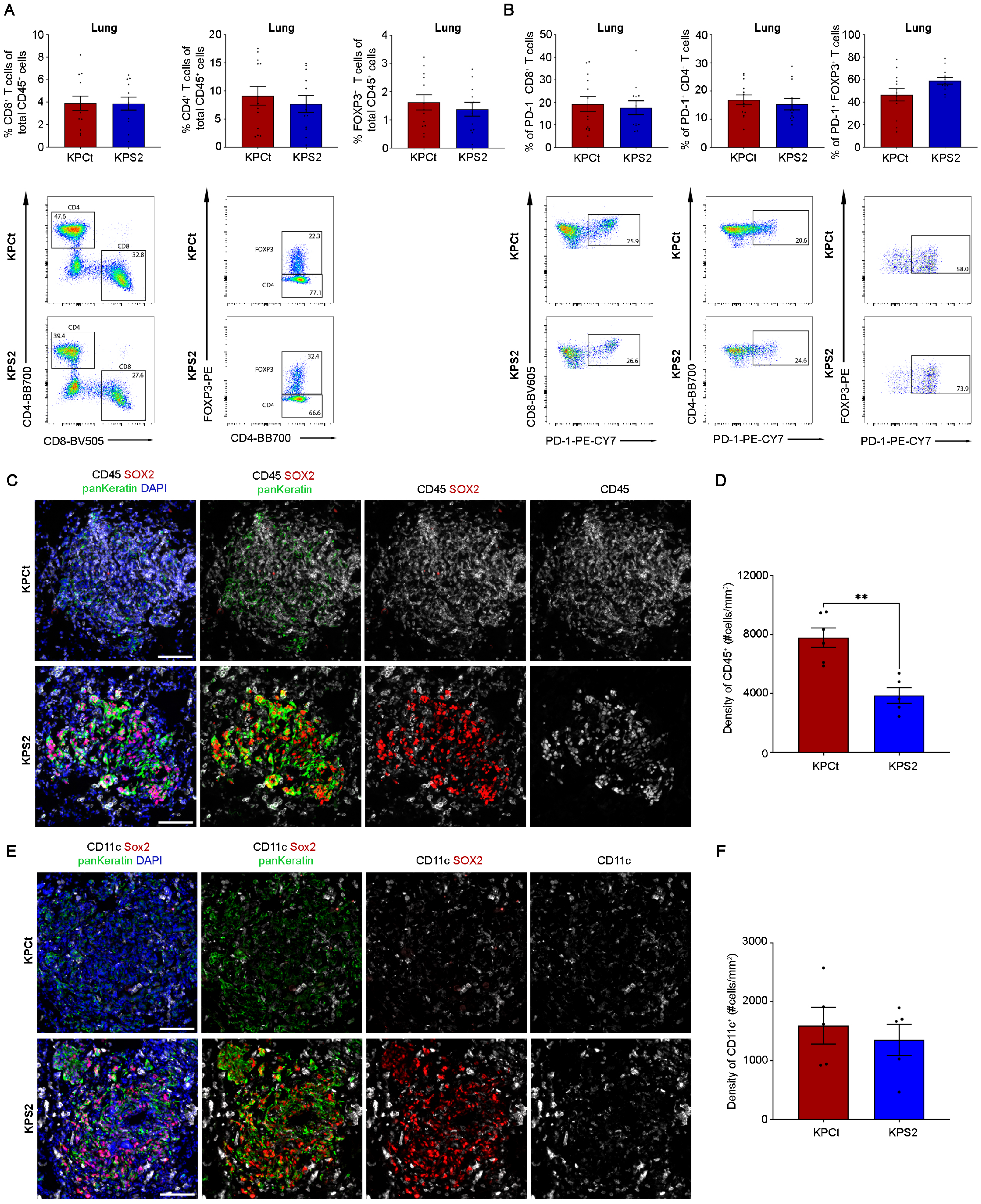

SUPPLEMENTARY FIGURE 4

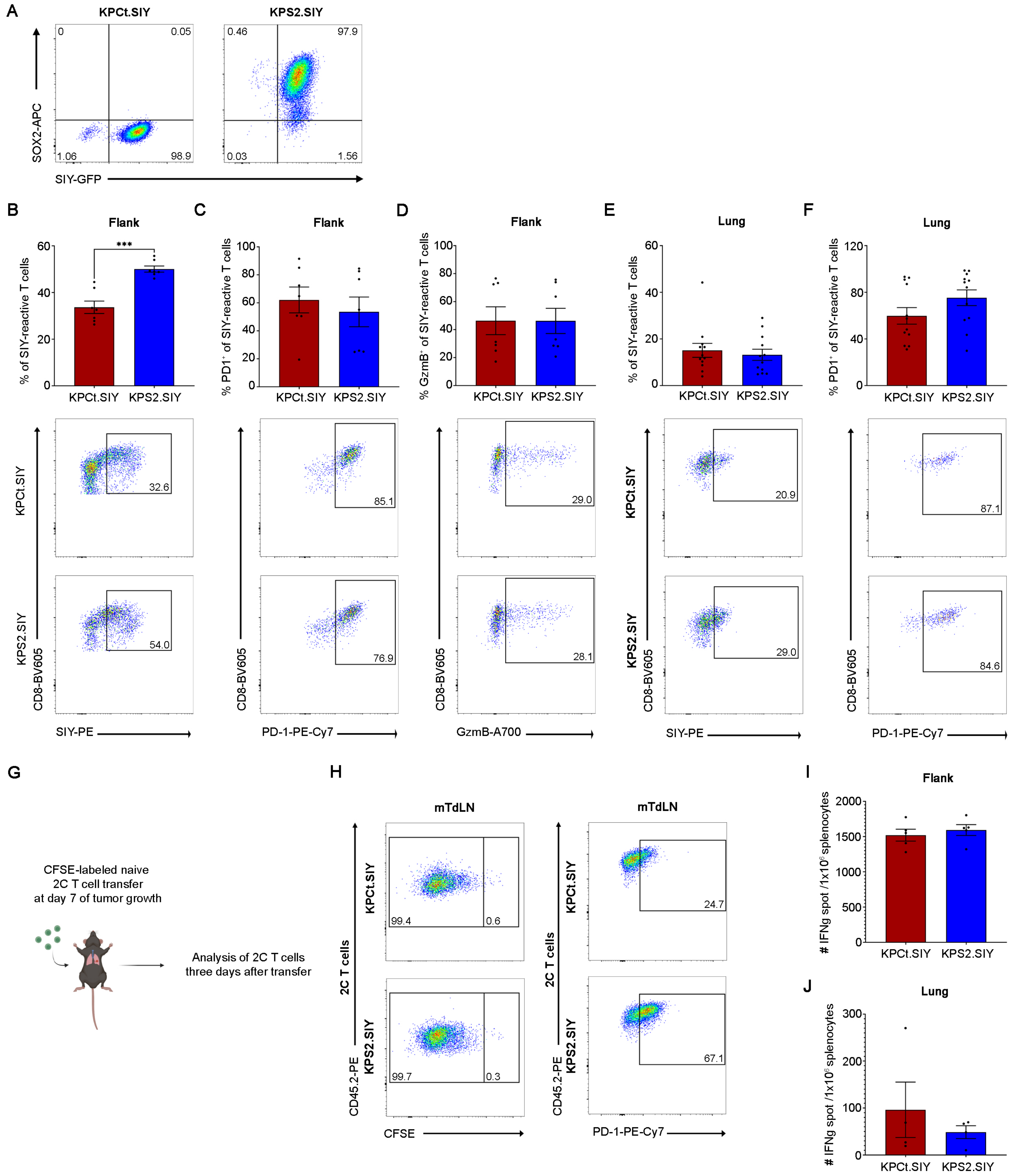

### SUPPLEMENTARY FIGURE 5

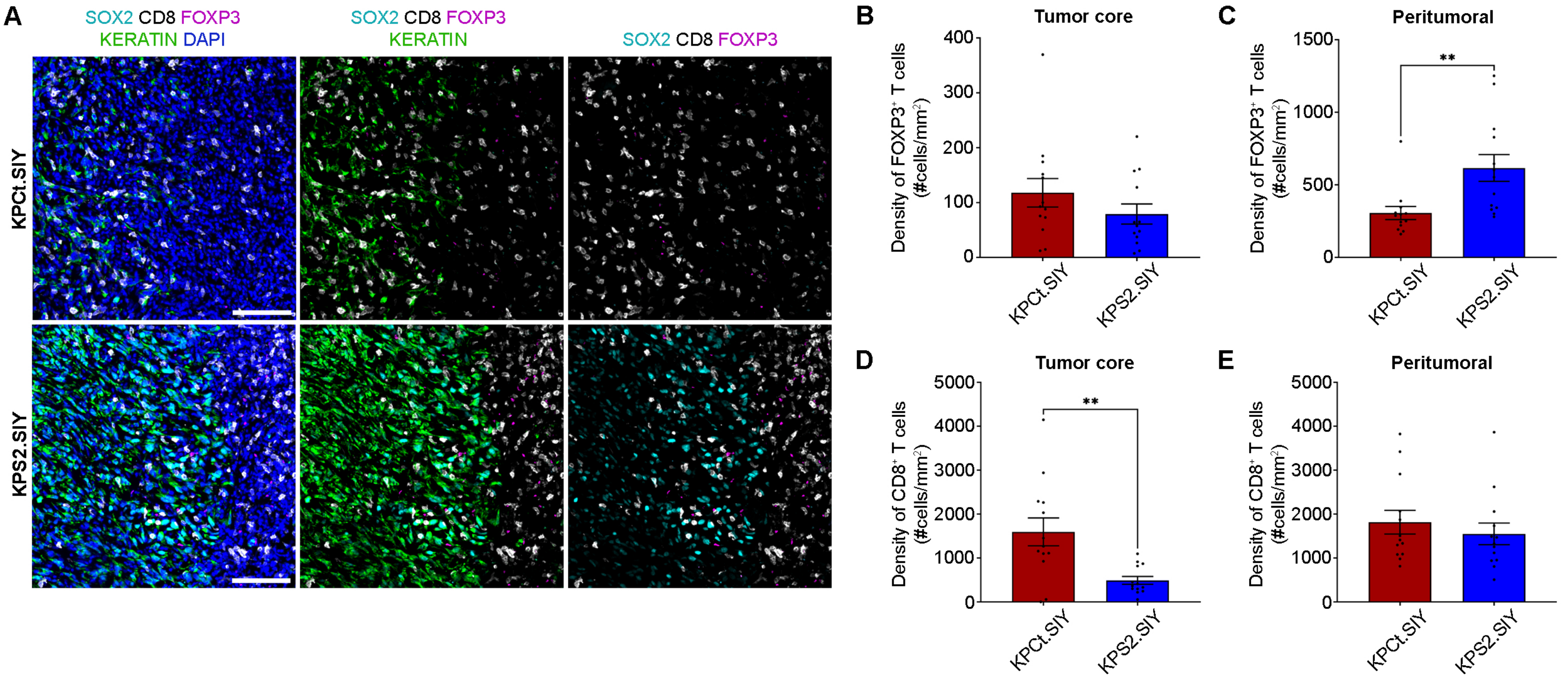

FIGURE SUPPLEMENTARY 6

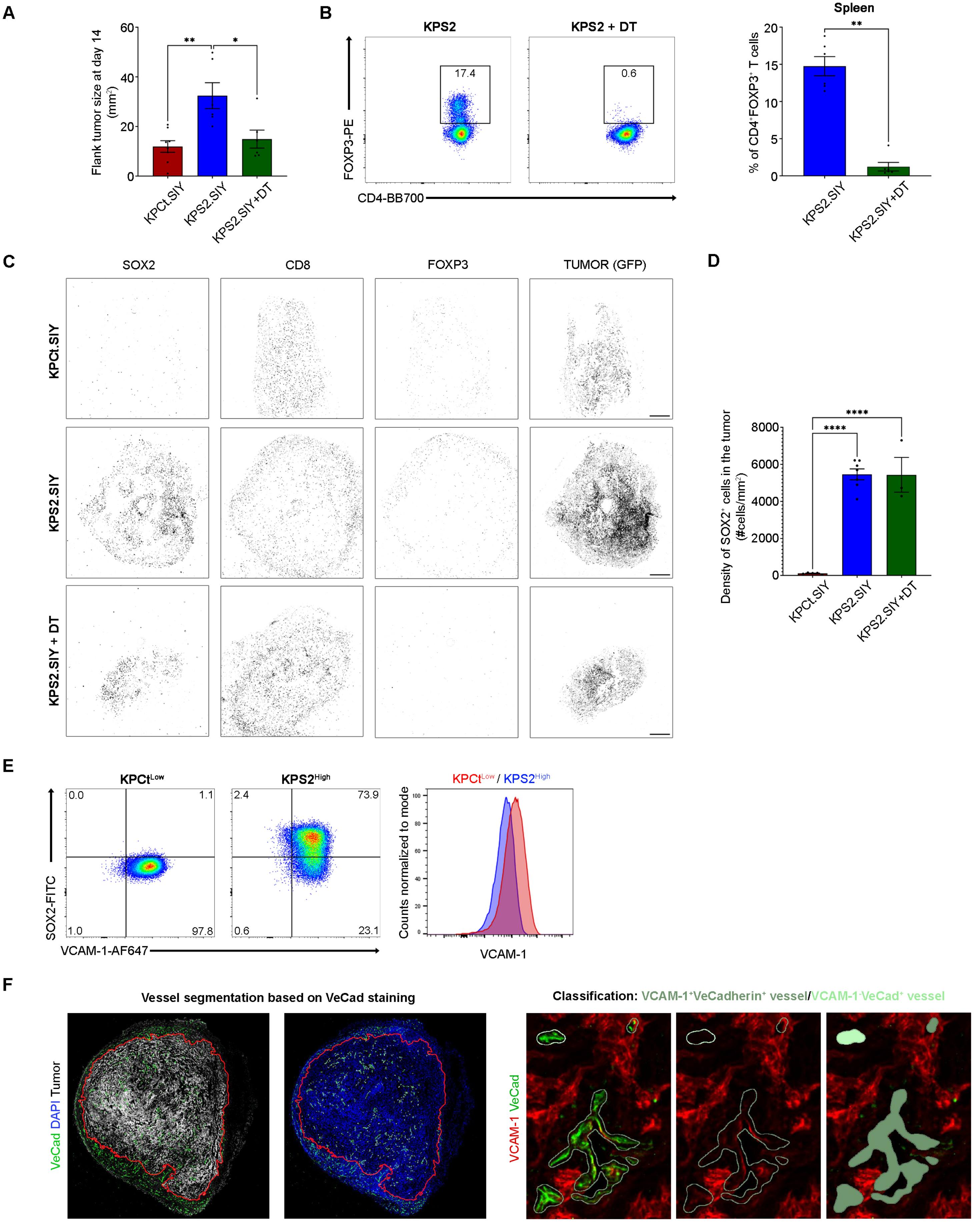

### SUPPLEMENTARY FIGURE 8

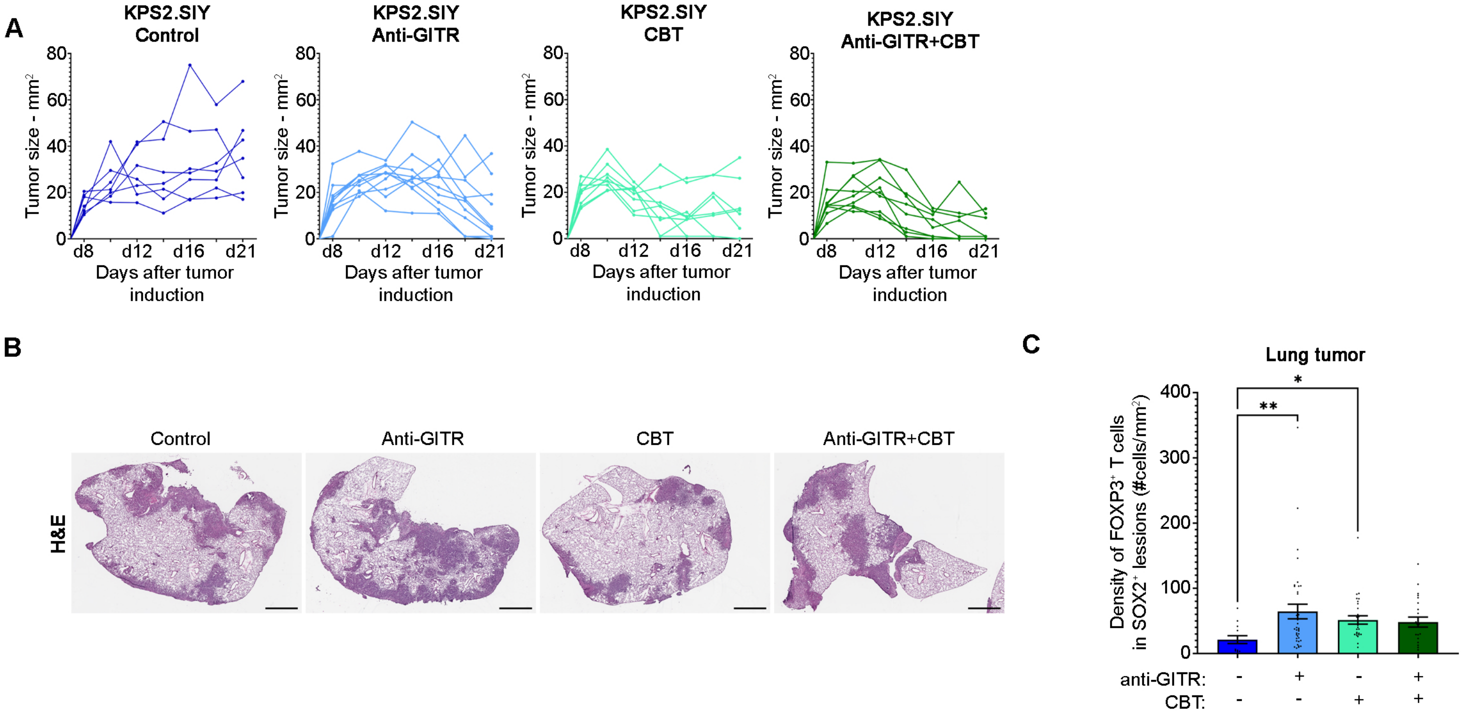
